# Metabolic signatures of regulation by phosphorylation and acetylation

**DOI:** 10.1101/838243

**Authors:** Kirk Smith, Fangzhou Shen, Ho Joon Lee, Sriram Chandrasekaran

## Abstract

Acetylation and phosphorylation are highly conserved post-translational modifications (PTMs) that regulate cellular metabolism, yet how metabolic control is shared between these PTMs is unknown. Here we analyze transcriptome, proteome, acetylome, and phosphoproteome datasets in *E.coli*, *S.cerevisiae*, and mammalian cells across diverse conditions using CAROM, a new approach that uses genome-scale metabolic networks and machine-learning to classify regulation by PTMs. We built a single machine-learning model that accurately distinguished reactions controlled by each PTM in a condition across all three organisms based on reaction attributes (AUC>0.8). Our model uncovered enzymes regulated by phosphorylation during a mammalian cell-cycle, which we validate using phosphoproteomics. Interpreting the machine-learning model using game-theory uncovered enzyme properties including network connectivity, essentiality, and condition-specific factors such as maximum flux that differentiate regulation by phosphorylation from acetylation. The conserved and predictable partitioning of metabolic regulation identified here between these PTMs can enable rational engineering of regulatory circuits.

**Graphical Abstract:** 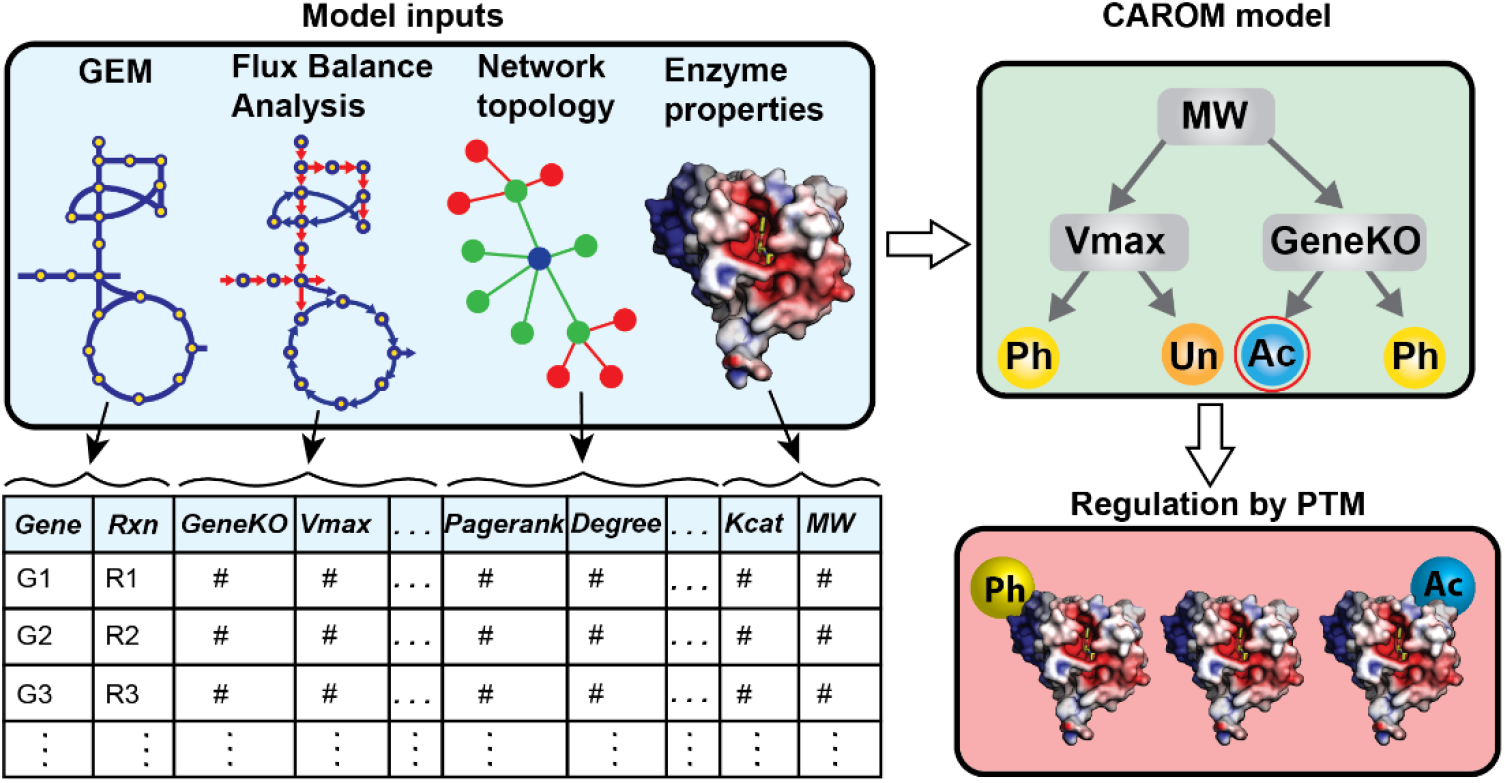

## Introduction

A key challenge in systems biology is to predict how various regulatory processes orchestrate cellular response to perturbations. Numerous mechanisms regulate metabolic response to new environments [1–8]. Nevertheless, it is unclear why or when some enzymes are regulated by acetylation while others through PTMs such as phosphorylation [3,4]. Several advantages of regulation by PTMs have been proposed over the past five decades [9–11]. These include low energy requirements, rapid response, and signal amplification. Yet these characteristics do not differentiate between PTMs such as acetylation and phosphorylation. The staggering complexity of each regulatory process has limited the comparative analysis of metabolic regulation at a systems level [3]. Existing studies have focused on a single regulatory process, usually transcriptional regulation [4,12–20]. Such studies have revealed reaction reversibility and metabolic network structure to be predictive of regulation [8,15,21–24]. Yet these studies do not shed light on the differences between each regulatory process, especially PTMs. In sum, although some general network principles of regulation are known, how it is partitioned among various regulatory mechanisms is unclear.

We hence developed a data-driven approach, called *Comparative Analysis of Regulators of Metabolism* (CAROM), to identify unique features of each PTM. CAROM achieves this by comparing various properties of metabolic enzymes, including essentiality, flux, molecular weight, and topology. It identifies properties that are more highly enriched among targets of each process than expected by chance. Using CAROM, we found features that were significantly associated with each PTM. Nevertheless, no single feature on its own is completely predictive of regulation. CAROM hence uses machine learning to uncover how features in combination influence regulation. We used CAROM to understand PTM dynamics during well-characterized fundamental processes in microbes and mammalian cells, namely the cell cycle, transition to stationary phase, and response to nutrient alterations. While we focus on acetylation and phosphorylation here as they are the most well-studied PTMs with available omics datasets, our approach can be applied to any regulatory process.

The manuscript is organized as follows: we first analyze various multi-omics datasets in *E. coli*, yeast and mammalian cells and reveal properties that are either enzyme-specific (molecular weight) or context-specific (flux) that correlate with regulation by each PTM. These common observations across various organisms allowed us to build a multi-organism machine-learning model that explains regulation in each condition using these features. The feature importance from CAROM is highly consistent across numerous studies in all organisms studied here. These results suggest that this approach is applicable to a wide range of model systems. CAROM can shed light on how metabolic changes impact PTMs. Proteomics surveys have found PTM sites on almost all metabolic enzymes [12,25]. A key challenge currently is the identification of condition-specific PTM sites and how they coordinately regulate metabolism in a condition [3,4,26]. Overall, CAROM provides a top-down, context-specific, enzyme property-based picture of metabolic regulation.

## Results

### Comparing regulation using CAROM

The CAROM approach takes as input a list of proteins that are the targets of one or more PTMs. CAROM analyzes the properties of the targets of PTMs in the context of a genome-scale metabolic network model. We hypothesize that target preferences of regulators can be inferred from the network topology and fluxes. CAROM compares the properties of the targets statistically using Analysis of Variance (ANOVA). It also builds a machine learning model capable of classifying regulation using boosted decision trees. Overall, CAROM compares the following 13 properties:

- Impact of gene knockout on biomass production, ATP synthesis, and viability across different conditions
- Flux through the network measured through Flux Variability Analysis, Parsimonious flux balance analysis (PFBA), and reaction reversibility
- Enzyme molecular weight and catalytic activity
- Topological properties, including the total pathways each reaction is involved in, its degree, betweenness, closeness, and PageRank

These properties were chosen based on ease of calculation using Flux Balance Analysis (FBA) and based on prior literature that have shown that hubs in the network and essential genes are frequent targets of transcriptional regulation [27]. Overall, CAROM can help interpret regulation in a condition and forecast targets of regulation using these features above. The CAROM source-code is available from the Synapse bioinformatics repository https://www.synapse.org/CAROM

### Shared features of enzymes regulated by acetylation and phosphorylation in yeast

We first analyzed the dynamics of metabolic regulation during a well-characterized process in yeast, namely, transition to stationary phase. We obtained RNA sequencing, time-course proteomics, acetylomics, and phospho-proteomics data from the literature [28–30]. Targets for each process were determined based on differential levels between stationary and exponential phase (Methods). We assumed that PTMs that are dynamic and conditionally regulated are likely to be functional [31].

Protein targets were mapped to corresponding metabolic reactions using the gene-protein-reaction annotations in the genome-scale metabolic network model of yeast [32]. There was significant overlap among reactions regulated through changes in both the transcriptome and proteome, and transcriptome and acetylome (hypergeometric p-value = 5 × 10^−25^ and 1 × 10^−15^ respectively, S. Table 1). In contrast, there was little overlap between targets of phosphorylation with other mechanisms (p-value > 0.1; S. Table 1). While prior studies found higher overlap between targets of PTMs [33,34], they used all possible sites that can be acetylated or phosphorylated. However, only a fraction of PTM sites are likely to be active and functional in a single condition. Overall, each regulatory mechanism had a distinct set of targets (Figure 1A). The targets of each regulatory mechanism were then used as input to CAROM.

**Figure 1.**
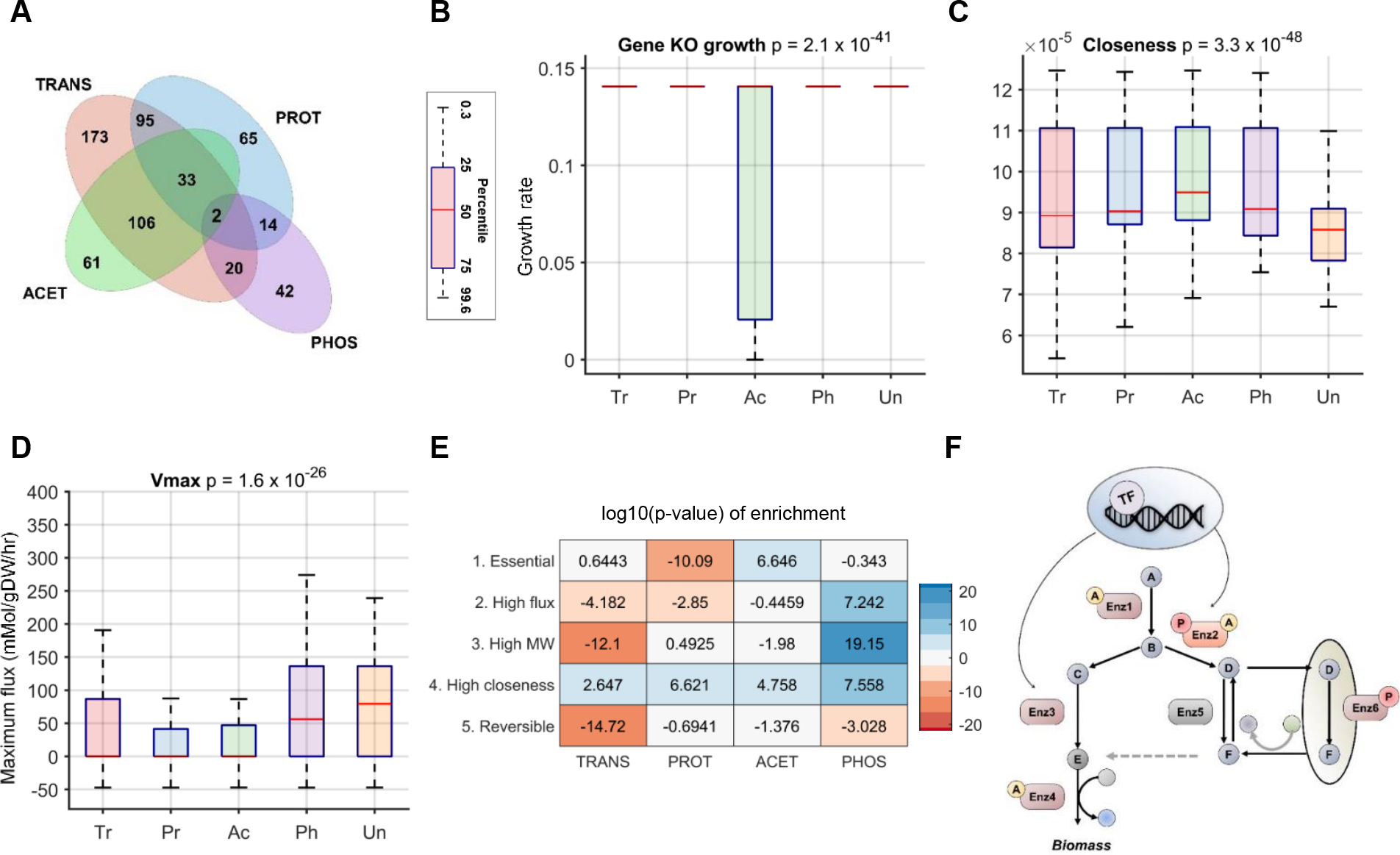
Comparison of the properties of the targets of regulation in yeast. The ANOVA p-value comparing the differences in means is shown in the title of the box plots. (Abbreviation: Enzymes regulated by transcription (Tr), post-transcription (Pr), acetylation (Ac), phosphorylation (Ph), Unregulated or unknown regulation (Un)) **A.** The Venn diagram shows the extent of overlap between targets of each process in stationary phase. Only 2 genes were found to be regulated by all four mechanisms. Targets of phosphorylation did not show any significant overlap with other mechanisms, while transcriptome and proteome showed the highest overlap (S. Table 1). **B.** Enzymes that impact growth when knocked out are highly likely to be acetylated. **C.** Enzymes with poor connectivity, as measured through the network connectivity metric - closeness, are more likely to be Unregulated. **D.** Enzymes catalyzing reactions with high maximum flux are likely to be either regulated through phosphorylation or to be unregulated. **E.** The heatmap shows the statistical enrichment (positive sign) and depletion (negative sign) of the targets of each process among reactions that are - (1) essential, (2) have high maximum flux (Vmax > 75^th^ percentile), (3) catalyzed by enzymes with high molecular weight (MW > 75^th^ percentile), (4) highly connected (Closeness > 75^th^ percentile), and (5) reversible. **F.** A schematic pathway summarizing the division of labor in metabolic regulation. Essential reactions (Enz1 and Enz4) are preferentially acetylated; reactions in futile cycles and in different compartments (Enz6) are phosphorylated, and reactions with high connectivity are regulated through multiple mechanisms (Enz2). Reversible reactions are predominantly unregulated or regulated by unknown mechanisms (Enz5).

We used CAROM to find common features of enzymes that are regulated by each mechanism. We first analyzed the regulation of enzymes that are essential for growth in minimal media. Essential enzymes in the yeast metabolic model were determined using FBA. Surprisingly, this set of enzymes was highly enriched among those regulated by acetylation but not by other processes (ANOVA p-value < 10^−16^; Figure 1B; S. Table 2). Since regulation can be optimized for fitness across multiple conditions [35], we identified enzymes that impact growth in 87 different nutrient conditions comprising various carbon and nitrogen sources using FBA. This set of essential enzymes was once again enriched for acetylation relative to other mechanisms (ANOVA p-value < 10^−16^; S. Figure 1). This trend was observed using an experimentally derived list of essential genes as well (hypergeometric p-value = 2 × 10^−7^ for acetylation). Thus, essential enzymes are likely to be constitutively expressed and their activity modulated through acetylation. This may explain why transcriptional regulation has minimal impact on fluxes in central metabolism, which contain several growth-limiting enzymes [3,14].

We next determined the impact of reaction position in the network on its regulation. We counted the number of pathways each reaction is involved in, along with other topological metrics, such as the closeness, degree, and Page Rank. We found that the regulation of enzymes differed significantly based on network topology (Figure 1C; S. Figure 2). First, reactions with low connectivity, measured through any of the topological metrics, were highly likely to be not regulated by these mechanisms. In contrast, highly connected enzymes linking multiple pathways were more likely to be regulated by PTMs. Connectivity metrics however were unable to differentiate between the two PTMs. Interestingly, reactions regulated by both PTMs had the highest connectivity (S. Figures 2, 3). Several key hubs, such as acetyl-CoA acetyltransferase, hexokinase and phosphofructokinase are regulated by multiple mechanisms (S. Table 3).

We next assessed how regulation differs based on the magnitude and direction of flux through the network. We inferred the full range of fluxes possible through each reaction using flux variability analysis (FVA) [36]. Since yeast cells may not optimize their metabolism for biomass synthesis during transition to stationary phase, we also performed FVA without assuming biomass maximization. We found that reversible reactions were not regulated by any of these mechanisms (S. Figure 4). A recent study found the same trend for allosteric regulation as well [21]. However, reversibility alone did not differentiate between regulatory mechanisms.

Interestingly, reactions that have high predicted maximum flux (Vmax) from FVA, such as ATP synthase and phosphofructokinase, were predominantly regulated by phosphorylation (Figure 1D; ANOVA p-value < 10^−16^). This set of phosphorylated reactions comprise several kinase-phosphatase pairs, enzymes that are part of loops that consume energy (“futile cycles”), or reactions that have isozymes in compartments such as vacuoles or nucleus (S. Table 4). Thus, phosphorylation in this condition selectively regulates reactions to avoid futile cycling between antagonizing reactions or those operating in different compartments. Using data from experimentally constrained fluxes from the Hackett *et al* study [21] revealed similar patterns of regulation (S. Figure 5).

Finally, we compared regulation based on fundamental enzyme properties: catalytic activity and molecular weight. While catalytic activity was similar across the targets of all mechanisms, targets of phosphorylation had the highest molecular weight (p-value < 10^−16^) (S. Figure 6). There is no correlation between molecular weight and maximum flux (Pearson’s correlation R = 0.02), suggesting that both maximum flux and molecular weight are likely to be independent predictors of regulation by phosphorylation.

To check if this pattern of regulation is observed in other conditions, we analyzed data from nitrogen starvation response and cell cycle in yeast, where both phospho-proteomics and transcriptomics data are available [37–40]. A similar trend of regulation was observed in this condition (S. Figure 6), with phosphorylation regulating isozymes and enzymes that have high Vmax (futile cycles). Overall, these results are robust to the thresholds used for finding differentially regulated sites, using data from different sources, and other modeling parameters (S. Tables 5, 6, 7, 8, 9).

### Context specific metabolic regulation by PTMs in *E. coli*

Since many mechanisms of metabolic regulation are evolutionarily conserved [3], we next analyzed multi-omic data from *E. coli* cells during stationary phase [41–43]. By analyzing transcriptomics, proteomics, acetylomics and phosphoproteomics data using the *E. coli* metabolic network model, we uncovered that the pattern of regulation observed in yeast was also observed in *E. coli* (Figure 2A-C, S. Figure 7). Essential reactions were enriched for regulation by acetylation, and reactions with high maximum flux or large enzyme molecular weight were enriched for regulation by phosphorylation. However, in contrast to yeast, phosphorylation impacted very few metabolic genes in *E. coli*, and may play a relatively minor role in this specific context. Phosphorylation had 20-fold fewer targets compared to other mechanisms, and its targets overlapped significantly with other processes (S. Tables 10, 11). Interestingly, the number of reactions with high maximum flux was considerably lower in *E. coli* compared to yeast (1282 in Yeast and 100 in E. coli), which correlates with the difference in phosphorylation between the species.

**Figure 2.**
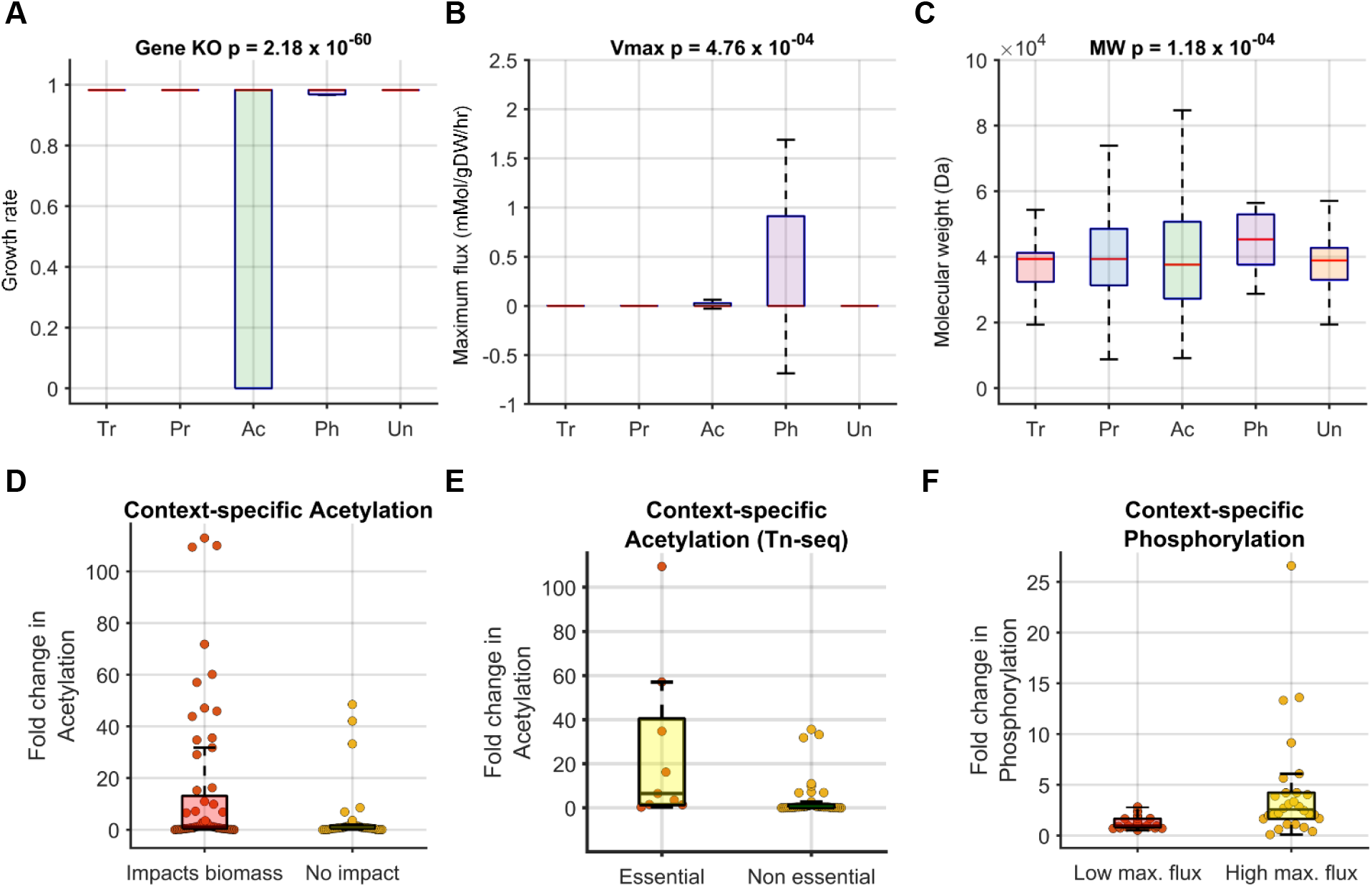
Comparison of the properties of enzymes in *E. coli* regulated by transcription (Tr), post-transcription (Pr), acetylation (Ac), phosphorylation (Ph) or Unregulated/Unknown regulation (Un) during transition to stationary phase. Similar to yeast, reaction essentiality **(A)**, maximum flux **(B)** and molecular weight **(C)** are predictive of regulation by acetylation and phosphorylation (Vmax, MW) respectively. Proteins that were found to be conditionally essential (growth < wild type glucose) based on FBA **(D)** or Transposon sequencing (Z-score < −2) **(E)** were more likely to be acetylated (p-value = 0.02 & 0.0011 for FBA and Tn-seq respectively). **F.** Enzymes that are predicted to have high maximal flux (Vmax > 90^th^ percentile) in a condition were likely to be phosphorylated compared to those with low maximal flux (p-value = 0.008).

Regulation by acetylation and phosphorylation are strongly associated with factors such as reaction flux and essentiality that change significantly between conditions. To further understand the condition-specific regulation of enzymes by PTMs, we used data from the Schmidt *et al* study that measured PTM levels for a small set of proteins in *E. coli* [44]. From this dataset we used 11 growth conditions in distinct nutrient sources that could be modeled using FBA. We selected 10 and 5 proteins, which were both part of the metabolic model and had acetylation and phosphorylation data, respectively. Despite the small sample size, we found that enzymes that impact biomass when deleted using FBA were more likely to be regulated by acetylation in that condition (p-value = 0.02; Figure 2D). This trend was also observed using experimental gene essentiality data from transposon mutagenesis screens (TN-seq) across these growth conditions (Figure 2E). For example, isocitrate lyase (aceA) show a consistent increase in acetylation as it becomes more essential (S. Figure 8, 9). Similarly, we observed a significant association between phosphorylation levels and the maximal flux through a reaction in each condition (Figure 2F). For example, phosphorylation of isocitrate dehydrogenase (icd) increased up to 20-fold in conditions with the highest maximal flux (S. Figure 10).

These results suggest that the metabolic features like essentiality and flux are predictive of both the regulation of different enzymes in a condition and for the same enzyme between conditions. Nevertheless, even though the maximal reaction flux and essentiality were associated with regulation by PTMs for many proteins in both organisms, there were exceptions that did not show this trend, suggesting that various factors identified earlier likely influence regulation by PTMs in a combinatorial fashion.

### Classifying metabolic regulation by PTMs using CAROM

While our statistical analysis has revealed the impact of various metabolic features on regulation by PTMs, each feature on its own is a weak predictor. We next sought to uncover how these features in combination determine the regulation of each enzyme. We used machine-learning (ML) to build a CAROM model that accounts for all these features and quantifies their interrelationship in influencing regulation by PTMs. While metabolic network models are more mechanistic, ML methods outperform metabolic models in prediction tasks [45]. Integrating metabolic network outputs with ML can enable mechanistic interpretation without compromising predictive accuracy [46,47]. We used the decision trees ML algorithm in CAROM due to its ease of interpretation and created an ensemble of decision trees using the XGBoost framework [48].

We re-analyzed the *E. coli* and yeast genome-wide omics datasets using CAROM. We further augmented this with phosphorylation and acetylation datasets from HeLa cells to assess if similar pattern of PTM regulation exists in mammalian cells. Time course acetylation data was taken from the Kori *et al* study [49], which identified 702 proteins whose acetylation levels changed significantly over time (Mann-Kendall test p-value < 0.05). Similarly, time course phosphorylation data from HeLa cells undergoing mitosis were obtained from Olsen *et al* [50].

We created a single CAROM model using data from all organisms with the goal of identifying conserved patterns of PTM regulation. A ternary classification algorithm was built to identify proteins that are regulated by acetylation, phosphorylation or were not regulated by these PTMs. The input to CAROM was the list of 13 features (Methods; Figure 3A, 3B). The model was trained using known examples of proteins that were regulated by each of the PTMs. The trained CAROM model was then used to predict the regulators of new proteins based on their feature values.

**Figure 3:**
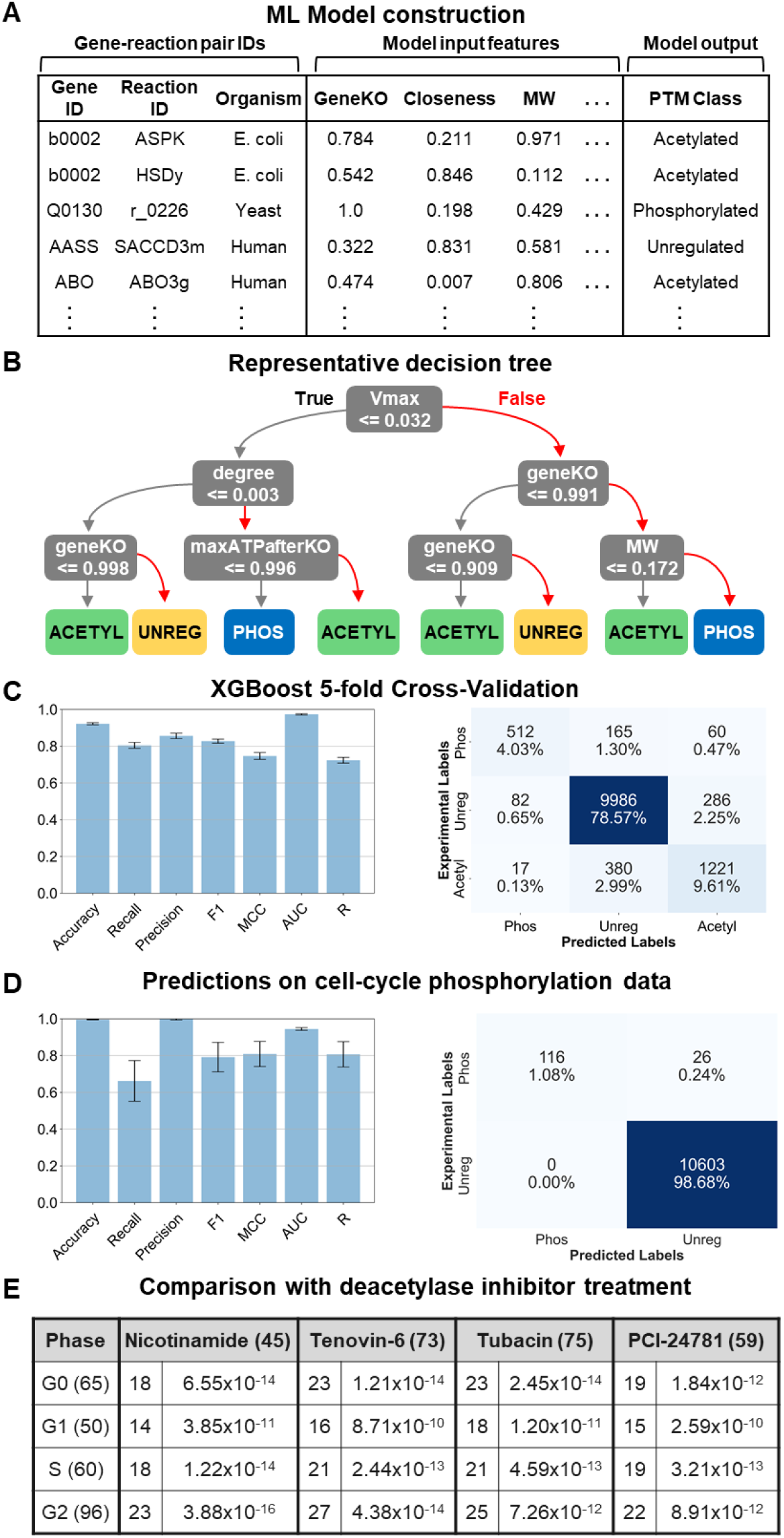
Construction and validation of the CAROM model. **A**. Table of inputs for CAROM. The input features comprise 13 gene, reaction, and enzyme properties. The target column includes the post-translational modification class. Each gene-reaction pair is marked as either phosphorylated, acetylated, or unregulated by PTMs. **B**. A single decision tree model was built by training on the observations from all organisms, while only using the top 50% most important features as identified in the SHAP analysis. The complexity of the tree was constrained by limiting the tree depth to enable ease of interpretation and visualization. The XGBoost model is made of an ensemble of such decision trees. **C**. The results from the CAROM model from 5-fold cross validation are shown in the bar graph (left) with the standard deviations represented by the error bars. The cross-validation results are also shown in the confusion matrix. **D.** Comparison of model predictions for the G1, S and G2 phases of the cell cycle with experimental phospho-proteomics data for those phases. Confusion matrix shows predictions from main CAROM model, while the bar graph shows the standard deviation for five models trained with different random seeds. **E**. Comparison of cell cycle acetylation predictions with experimental acetylomics data from HeLa cells treated with pan-deacetylase inhibitors. The number of unique acetylated genes for each group are displayed in parentheses. Within the table, the number of overlapping genes between each phase and drug is shown, along with the p-value of the hypergeometric test.

The trained CAROM model showed very high accuracy in predicting proteins that are regulated by each PTM in all three systems based on five-fold cross-validation, wherein a portion of the dataset (20%) is hidden from the model. We used a range of metrics to quantify accuracy including the Matthews Correlation Coefficient (MCC), the F1 score, precision, and recall. The ML models performed accurately based on all these metrics and significantly better than random shuffling of the data (Figure 3C).

To test the generalizability of this approach in novel conditions, we used the model to predict phosphorylation during a mammalian cell cycle. We used time-course phosphoproteome data for the first cell cycle from a murine lymphocyte cell line in response to a cytokine activation (Methods). We focused on the cell cycle as it is a fundamental process and is known to involve coordination of kinases and phosphorylase cascades [51]. Importantly, this model system was previously used by Lee *et al* to measure metabolomics changes during the cell cycle [52]. Phospho-proteomes were obtained at the same time points as the metabolomics data from the Lee *et al* study. We used the extracellular and intracellular metabolomics data from the Lee *et al* study to build metabolic models for each phase of the cell cycle. We used the DFA approach, a variation of dynamic FBA, to fit the rate of change of metabolites in FBA to experimental measurements from time course metabolomics [53,54]. We used this approach to create four different models corresponding to different phases of the cell cycle (G0, G1, G1-S and G2/M) (S. Figure 11, Methods).

The feature data (i.e., fluxes, topology) from the phase-specific metabolic models were used as input for the CAROM model to predict reactions regulated by phosphorylation. The G0 phase data was used for additional training of the model to learn cell-type specific phosphorylation patterns, and the G1, G2 and S phase were used for testing the CAROM model. CAROM achieved high MCC, AUC and precision in all conditions tested. 116 out of 142 predictions on phase-specific phosphorylated enzymes/reactions were also observed experimentally (S. Table 12). Similar to *E. coli* and yeast, there was significant correlation between the maximum flux of a reaction in a condition and the change in phosphorylation of the corresponding enzyme during the mammalian cell cycle (S. Figure 11). For example, AMP deaminase (AMPD2) shows a threefold increase in phosphorylation in G2 phase wherein it also shows a corresponding increase in maximal flux. These results together suggest that knowledge of fluxes can be predictive of regulation by phosphorylation in mammalian systems as well.

CAROM also predicted several reactions to be targets of acetylation in various phases (S. Table 13). The predicted list includes enzymes such as ATP-citrate lyase whose activity is known to be regulated by acetylation during the cell cycle [55,56]. As we lack proteome-wide time-course acetylation data to systematically confirm these predictions, we compared predictions with data from cells treated with deacetylase inhibitors [57]. Deacetylase inhibitors prevent the removal of acetylation marks. Hence new acetylation marks progressively accumulate over time resulting in cell death. We hypothesized that acetylation sites predicted by the CAROM model during the cell cycle will be enriched among the proteins with increased acetylation after deacetylase inhibitor treatment. Indeed, there was a significant overlap between CAROM predicted acetylated enzymes and those found to increase significantly (> 1.5-fold) after treatment with four different pan-deacetylase inhibitors – nicotinamide, tenovin-6, tubacin and PCI24781. Interestingly, even though the experimental proteomics data was not phase specific, we observed the highest overlap for nicotinamide targets with CAROM predictions in the G2 phase of the cell cycle (hyper-geometric p-value = 3 × 10^−16^), which also had the highest number of acetylated reactions (Figure 3E; S. Table 14). This overlap suggests that growth inhibition likely occurs in the G2 phase, which is consistent with experimental data from nicotinamide treatment in various mammalian cell types that have observed growth arrest at G2 [58–60].

### Interpreting the machine-learning model using Shapley analysis

To understand how CAROM predicted regulation by each PTM, we used a game-theoretic framework called Shapley analysis to quantify the contribution of each feature to the model accuracy using the SHAP (SHapley Additive exPlanation) Python package [61,62]. The Shapley ‘feature importance’ values are computed by sequentially adding one feature at a time and measuring the feature’s contribution to the model output. To account for the order in which the features are added to the decision trees, this process is repeated for all possible orderings. The Shapley value represents the average impact for each feature across all orders (Methods).

All 13 features contributed to the CAROM predictions, albeit to various extents. Molecular weight and maximum flux had two of the highest importance scores, and higher values favored phosphorylation, which is consistent with the high enrichment we observed using our statistical analysis (Figure 4A). Growth-related features, such as impact of gene knockout on biomass and ATP, were found to have opposite Shapley values for acetylation and phosphorylation respectively (Figure 4A). Thus, high growth values after knockout favor phosphorylation while low growth values favor acetylation. Similar to *E. coli* and yeast, the set of proteins acetylated in HeLa cells were highly enriched for essential genes identified by both FBA simulations and experimental genome-wide CRISPR knockdown studies (hypergeometric test comparing acetylated metabolic genes to all metabolic genes, p-value = 1 × 10^−3^ & 9 × 10^−7^ for FBA and CRISPR respectively). These results show that changes in fluxes and essentiality between conditions are associated with a corresponding change in regulation by PTMs.

**Figure 4:**
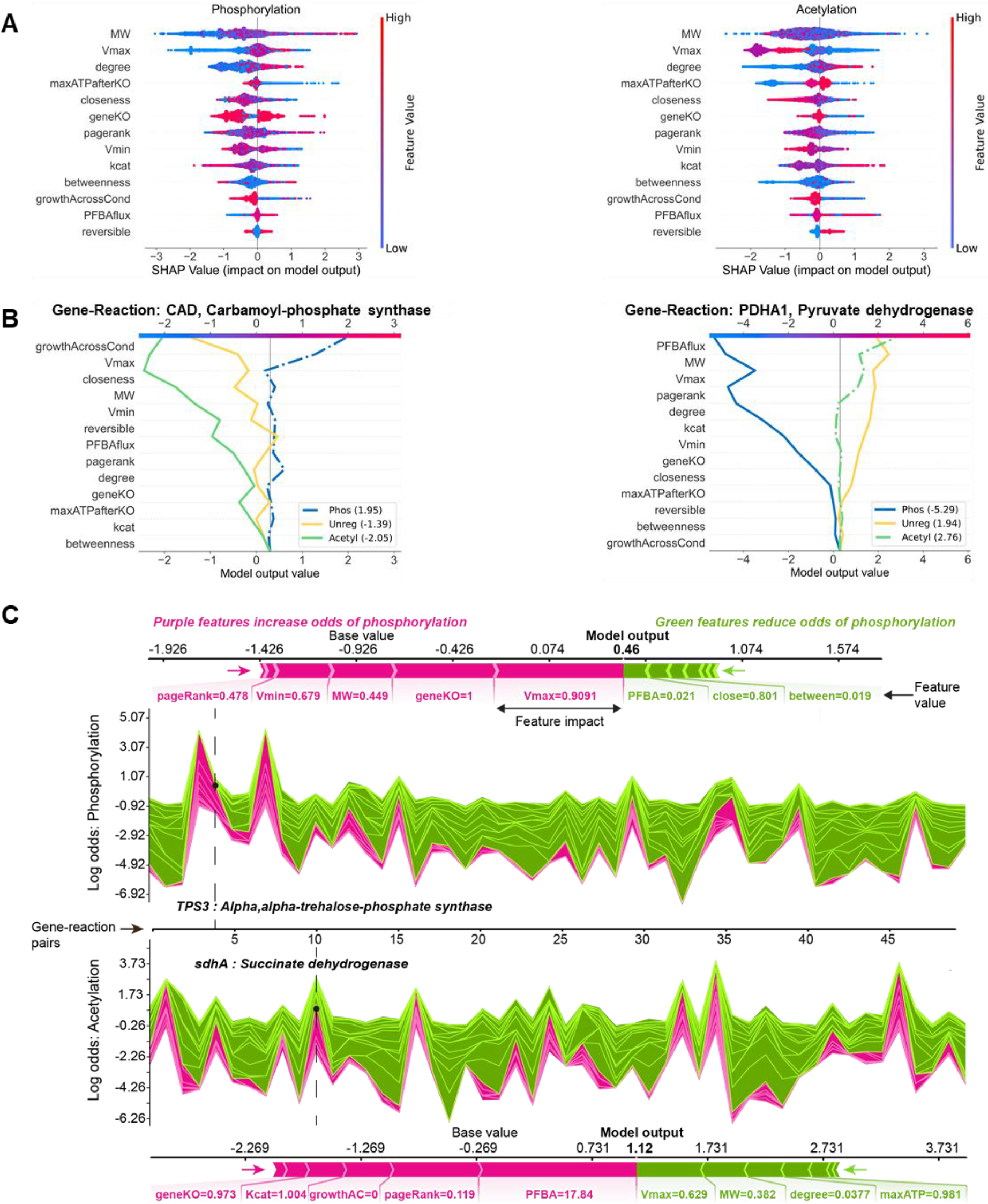
Interpretation of the CAROM model using Shapley analysis. **A.** SHAP summary plot for the phosphorylation class (left) and acetylation class (right). The summary plot shows how a feature’s effect on the output changes with its own value. For each feature, high values are shown in red and low values in blue. For example, it appears that Vmax is positively and negatively correlated with the log odds of phosphorylation and acetylation, respectively. Features are ordered on the y-axis by their average SHAP importance value across the three classes. **B.** SHAP decision plots for a phosphorylated enzyme (left) and acetylated enzyme (right) show how the model’s prediction was made for a single observation. Each line represents the log odds for a single class. The features are on the y-axis and are sorted by the average SHAP value for that specific observation. The lines intercept the top x-axis at their final log odds value. The class with the maximum log odds value is used as the model’s output. **C.** SHAP force plots show the features which significantly pushed the model output from its expected value to its final prediction. Features that push the prediction higher for the respective class are shown in purple and features that pushed it lower are shown in green. Single force plots for a phosphorylated reaction (top; TPS3) and an acetylated reaction (bottom; sdhA) are shown. The collective force plots are made up of many single force plots rotated 90 degrees and stacked together horizontally and are shown for phosphorylation (upper middle) and acetylation (bottom middle) for the same 50 random observations. The model output, f(x), is on the y-axis and observations on the x-axis. The dashed lines show where the single force plot observations appear in the collective force plot. For both the single and collective force plots, the model output is read where the purple and green areas intersect.

Molecular weight, topological features and reversibility were used by CAROM to differentiate all regulated genes from those that are un-regulated (Figure 4A, 3B, S. Figure 12). Gene knockout growth and maximum flux likely aid in differentiating between PTMs based on their opposing Shapley values for each PTM. These observations help explain why using both acetylation and phosphorylation in a single model improves performance compared to ML models built separately for each PTM (S. Figure 14). The SHAP decision plots and force plots shows how these features influence the prediction outcome for any given protein (Figure 4B). This also allowed us to identify factors that led to incorrect predictions by the ML model. Notably, a majority of the incorrect phosphorylation predictions were on proteins that had high molecular weight (S. Figure 13). Our ability to more accurately predict context specific fluxes and gene essentiality in the future may help rectify these incorrect predictions.

To tease out organism specific differences, we next built CAROM models separately for each organism. Overall, the model accuracy and feature importance were similar for both the pan-organism CAROM model and organism-specific models (S. Figures 15, 16, 17, 18). This suggests that a similar template involving the same set of features is used for partitioning regulation. Vmax, molecular weight, topology and gene knockout values are used in the same way in all three organisms for partitioning regulation. However, the specific parameters (the threshold for Vmax or molecular weight) were organism specific. Nevertheless, these parameters can be learned by CAROM using a small subset of data. Hence while the accuracy is very low when an entire organism’s data is removed from the model and used as a test set, a substantial increase is observed when just 10% of the test organism’s data is used for additional training (S. Figure 18).

The distribution of these top features from CAROM may explain the differences in distribution of PTMs observed between different species and metabolic conditions. We observed that the number of reactions with high Vmax was an order of magnitude higher in yeast compared to *E. coli* for the same condition (stationary phase). A concordant difference in number of reactions regulated by phosphorylation was observed between the two species (S. Figure 19). A similar trend was observed in phosphorylation levels in different conditions within the same species, namely the phases of the mammalian cell cycle and nutrient adaptation in E. coli (S. Figures 10,11). In addition, the total reactions regulated by acetylation correlated with the number of growth-limiting enzymes across conditions or species (S. Figure 8, 9, 19).

## Discussion

There are several ways to regulate an enzyme’s activity in a cell. Yet, the principles that determine when an enzyme is regulated by different PTMs are unknown. Here we systematically analyze patterns of metabolic regulation in model microbes and mammalian cells using a new approach called CAROM. Our approach explains why some proteins are regulated by specific PTMs in a given condition based on their biochemical properties, activity in a condition, and location in the metabolic network. We find that a small set of 13 features can distinguish the targets of each mechanism. The importance of these features is highly consistent across numerous datasets suggesting that these features may play a role in influencing regulation. Although the relevance of some of the features, such as topology, has been observed previously for transcriptional regulation, this is the first time that an association between regulation by PTMs and condition-specific attributes such as maximal flux has been reported.

These key features identified by CAROM may be related to specific functions performed by each PTM. For example, phosphorylation may represent a mechanism of feedback regulation to control futile cycles and high flux reactions that consume ATP [6,63]. The differences in the total number of isozymes and high flux enzymes between species may explain the varying number of phosphorylation targets observed between the species. Since isozymes arise frequently from gene duplication, our results may also explain the observation that duplicated genes are more likely to be regulated by phosphorylation [64]. However, it is unclear how the maximum flux is sensed by cells. These regulatory interactions may have been shaped by evolution to avoid drain of ATP. Cells may also utilize ‘flux sensors’ to identify such reactions [65]. Similarly, we find that enzymes are likely to be acetylated in conditions where their activity is growth limiting. The number of acetylated enzymes correlates with the number of essential genes between organisms or between conditions. During transition to stationary phase, essential genes do not show significant changes in transcript and protein levels, but show significant changes in acetylation in both yeast and *E. coli*. By regulating growth limiting enzymes, acetylation may play an evolutionarily conserved role in determining the balance of biosynthetic and catabolic processes in a cell.

Our approach does have limitations primarily due to the underlying algorithms and datasets used. The accuracy of the metabolic reconstruction strongly influences CAROM accuracy. False positive gene knockout essentiality predictions can lead to incorrect assignment of regulation by acetylation. Using experimental gene deletion screens can improve accuracy but may not be available for all conditions. Similarly, phosphorylation predictions can be impacted by flux predictions by FBA. FBA is currently the most powerful approach to obtain genome-wide fluxes. Nevertheless, the incorporation of context-specific omics datasets can improve accuracy of the predicted fluxes from FBA and subsequently predicted regulation by CAROM. Further, the set of features used in CAROM, although most of them were significantly associated with regulation, are unlikely to be exhaustive. These features were selected based on prior knowledge and ease of prediction using FBA. Other features such as presence of other PTMs may provide additional information to improve accuracy. Finally, ML methods require numerous measurements for training and may not perform well in cases with small sample sizes.

In sum, our analysis reveals a unique distribution of regulation by PTMs within the metabolic network. This can help identify PTMs that will likely orchestrate flux adjustments based on reaction attributes. By identifying context-specific factors that are associated with regulation by PTMs, CAROM can complement sequence-based approaches for identifying PTM sites. It is well established that individual regulators such as transcription factors or kinases have their own unique set of targets. Here we find that similar specialization likely occurs at a higher scale, between PTMs. Our approach can guide drug discovery and metabolic engineering efforts by identifying regulators that are dominant in different parts of the network [66]. CAROM can also be used to uncover the impact of metabolic alterations on PTMs in normal and pathological processes. Given the conservation of these principles in *E. coli*, yeast, and mammalian cells, it provides a path towards a detailed understanding of post-translational regulation in a wide range of organisms and to uncover target specificities of other PTMs. This approach may help define the basic regulatory architecture of metabolic networks.

## Methods

### Compilation of omics data

We used RNA-sequencing data from Treu *et al* 2014 that compared the expression profile of *S. cerevisiae* between mid-exponential growth phase with early stationary phase [30]. A 2-fold change threshold was used to identify differentially expressed genes. Lysine acetylation and protein phosphorylation data were obtained from the Weinert *et al* 2014 study that compared PTM levels between exponentially growing and stationary phase cells using *stable isotope labeling with amino acids in cell culture* (SILAC) [29]. A 2-fold change threshold of the protein-normalized PTM data was used to identify differentially expressed PTMs. Proteomics data was taken from Murphy *et al* time-course proteomics study [28]. The hoteling T2 statistic defined by the authors was used to identify proteins differentially expressed during diauxic shift; the top 25% of the differentially expressed proteins were assumed to be regulated. Proteomics data from Weinert *et al* was also used as an additional control and we observed the same trends using this data as well (S. Table 7). Further, we repeated the analysis after removing genes that were not expressed during transition to stationary phase; the transcripts for a total of 12 genes out of the 910 in the model were not detected by RNA-sequencing in the Treu *et al* study [30]. Removing the 12 genes did not impact any of the results (S. Table 6).

As additional validation, we used periodic data from the yeast cell cycle. Time-course SILAC phospho-proteomics data was obtained from Touati *et al* [39]. Phospho-sites whose abundance declined to less than 50% or increased by more than 50% at least two consecutive timepoints were considered dephosphorylated or phosphorylated respectively as defined by the authors. Transcriptomics data was taken from Kelliher *et al* study that identified 1246 periodic transcripts using periodicity-ranking algorithms [40]. The phospho-proteomics and transcriptome data during nitrogen shift was obtained from Oliveira *et al* [37,38]. The nitrogen shift studies compared the impact of adding glutamine to yeast cells growing on a poor nitrogen source (proline alone or glutamine depletion) with cells growing on a rich nitrogen source (glutamine plus proline). A 2-fold change threshold was used to identify differentially expressed transcripts and phospho-sites.

*E. coli* acetylation data was taken from the Weinert *et al* study comparing actively growing exponential phase cells to stationary phase cells [43]. Proteomics and transcriptomics were from Houser *et al* study of *E. coli* cells in early exponential phase and stationary phase [42]. Phospho-proteomics data for exponential and early stationary phase *E. coli* cells was taken form Soares *et al* [41]. We used a 2-fold change (p < 0.05) threshold for all studies.

Condition specific PTM data for *E. coli* was taken from Schmidt *et al* 2016 study [44]. Among the 22 different experimental conditions measured, those conditions that involved change in carbon sources that could be modeled using FBA were chosen. The following carbon sources were used: acetate, fumarate, galactose, glucose, glucosamine, glycerol, pyruvate, succinate, fructose, mannose and xylose. Out of 44 unique lysine acetylation and 21 serine/ threonine phosphorylation sites identified in the study (FDR < 0.01), 11 and 5 proteins were mapped to the metabolic model for the subset of conditions analyzed here. Protein modifications were normalized by their corresponding protein levels.

Acetylated proteins in HeLa cells were taken from Kori *et al* 2017 which measured time course acetylation levels in HeLa cells grown on 13C labeled glucose with samples collected at 0.5, 1, 4, 8, 12, 16, and 24 hours [49]. A total of 702 unique target proteins were identified based on significance of acetylation incorporation as monotonic trend across the time points using the Mann-Kendall statistical test (p-value < 0.05) as defined by the authors. For the phosphorylation data for HeLa cells, phosphorylation sites that are up-regulated during mitosis and show more than 50% occupancy as defined by the authors were used [50].

Phosphoproteomics data from the mammalian cell cycle contained a total of 5861 identified phosphopeptides. Phospho-peptides whose abundance intensities (or signal to noise ratios) are zero at any channel (or any time point sample), those with Ascore < 13, and those that were identified by a decoy dataset in a reverse manner were removed, resulting in a set of 3095 phosphopeptides that correspond to 1552 unique proteins. A z-score normalization was performed to identify phase specific differential levels of phosphorylated proteins (z threshold of +/− 2)

Gene essentiality based on CRISPR knockout screens was obtained from Hart *et al* 2015 study that measured essentiality across all 5 cell lines (HeLa, RPE1 DLD1, GBM and HCT116) [67]. Growth limiting genes with FDR < 0.05 were considered to be essential, as defined by the authors. In addition, essential genes from Hart *et al* 2017 study using genome-wide knockout screens in 17 human cell lines also showed similar enrichment among acetylated proteins (p-value = 1.7 × 10^−7^) [68].

The results are robust to the thresholds used for identifying differentially expressed genes or proteins (S. Tables 6, 7, 8). In all studies, genes and proteins that are either up or down regulated were considered to be regulated. The final data set table used for all comparative analyses is provided as a supplementary material (S. Tables 14, 15, 16).

### Genome scale metabolic modeling

We used the yeast metabolic network reconstruction (Yeast 7) by Aung *et al*, which contains 3,498 reactions, 910 genes and 2,220 metabolites [32]. The analysis of *E. coli* data was done using the IJO1366 metabolic model [69] and the mammalian cell cycle modeling was done using the human metabolic reconstruction (Recon1) [70]. All analyses were performed using the COBRA toolbox for MATLAB [71].

The impact of gene knockouts on growth was determined using flux balance analysis (FBA). FBA identifies an optimal flux through the metabolic network that maximizes an objective, usually the production of biomass. A minimal glucose media (default condition) was used to determine the impact of gene knockouts. Further, gene knockout analysis was repeated in different minimal nutrient conditions to identify genes that impact growth across diverse conditions; these conditions span all carbon and nitrogen sources that can support growth in the metabolic models. The number of times each gene was found to be lethal (growth < 0.01 units) across all conditions was used as a metric of essentiality.

To infer topological properties, a reaction adjacency matrix was created by connecting reactions that share metabolites. We used the Centrality toolbox function in MATLAB to infer all network topological attributes including centrality, degree and PageRank. Removing highly connected metabolites did not affect the associations between topology and regulation (S. Figure 20).

Flux Variability Analysis (FVA) was used to infer the range of fluxes possible through every reaction in the network. Two sets of flux ranges were obtained with FVA – the first with optimal biomass and the latter without assuming optimality. In the second case, the fluxes are limited by the availability of nutrients and energetics alone, thus it reflects the full range of metabolic activity possible in a cell. Reactions with maximal flux above 900 units were assumed to be unconstrained and were excluded from the analysis, as they are likely due to thermodynamically infeasible internal cycles [72]; the choice of this threshold for flagging unconstrained reactions did not impact the distribution between regulators over a wide range of values (S. Table 9).

For fitting experimentally derived flux data from Hackett *et al* [21], reactions were fit to the fluxes using linear optimization and the flux through remaining reactions that do not have experimentally derived flux data were inferred using FVA. Analysis using a related approach for inferring fluxes – PFBA, did not reveal any significant difference as PFBA eliminates futile cycles and redundancy by minimizing total flux through the network while maximizing for biomass [73] (S. Figure 5).

Reaction reversibility was determined directly from the model annotations. We also used additional reversibility annotation from Martinez *et al* based on thermodynamics analysis of the Yeast metabolic model [74]. Pathway annotations and enzyme molecular weight values were obtained from Sanchez *et al*. The catalytic activity values were obtained from Sanchez *et al*, Heckman *et al*, and Yeo *et al* for Yeast, *E. coli* and mammalian cells respectively [75–77]. The comparative analysis of regulatory mechanisms was also repeated using the updated Yeast 7.6 model and yielded similar results (S. Table 5) [75].

Models for each cell cycle phase were built using the Dynamic Flux Activity (DFA) approach [53,78]. The cell cycle metabolomics data contains 155 intracellular metabolites and 173 extracellular metabolites and was used as inputs for DFA. The time points were grouped in to different phases as follows: 0 – 4 hours for G0-G1, 4 – 8 – 12 hours for G1, 12 – 16 hours for G1-S, and 16 – 20 hours for G2-M. DFA utilizes time-course metabolomics data and calculates the rate of change of each metabolite level over time (d***M***/dt). The rate of change of each metabolite is calculated using linear regression in DFA. Based on the regression line for a metabolite *i*, one calculates *ϵ_i_* which is the slope divided by the intercept which is a normalization factor at the initial time point. Then, together with a known metabolic network for the stoichiometry matrix, ***S***, and by introducing flux activity coefficients, ***α*** and ***β***, the DFA equation becomes a modified version of the conventional FBA: ***S•v*** + ***α*** - ***β*** = ***ϵ***. ***α*** and ***β*** are both positive values. This equation is then solved by minimizing ***α*** + ***β*** and maximizing the biomass objective function, yielding a flux vector or distribution of all reactions for time-course data. For validating the CAROM model, the fluxes from the G0 phase were used in the training set and the remaining phases were used for testing. This analysis was repeated by training on different phases of the cell cycle. The accuracy from the G1, S and G2 phases was lower compared to training on G0. suggesting that these conditions have a distinct phosphorylation pattern from the G0 condition (S. Figure 21).

The comparative analysis of target properties was done using gene-reaction pairs rather than genes or reactions alone. The gene-reaction pairs accounts for regulation involving all possible combinations of genes and associated reaction. This includes isozymes that may involve different genes but the same reaction, or multi-functional enzymes involving same the gene associated with different reactions. For example, the 910 genes and 2310 gene-associated reactions resulted in 3375 unique gene-reaction pairs in yeast.

### Statistical analysis

All statistical tests were performed using MATLAB. Significance of overlap between lists was estimated using the hypergeometric test. Significance of the differences in target properties between regulatory mechanisms were determined using ANOVA, the non-parametric Kruskal-Wallis test, and after multiple hypothesis correction (S. Table 5).

### Machine learning

The CAROM-ML model was built using the XGBoost package in Python. XGBoost is a gradient boosting algorithm that uses decision trees as its weak learners [48]. Unlike bagging algorithms, such as random forest, which train their learners independently in parallel, boosting algorithms train their predictors sequentially. Each weak learner uses gradient descent to minimize the error of the previous learner. XGBoost is unique among boosted algorithms due to its speed and regularization abilities, which help prevent over-fitting.

We used a randomized search with an internal cross validation in the training set to tune hyperparameters. A stratified split was employed to ensure the class balance was preserved between the training and test sets. To measure the model robustness and generalization, we performed 5-fold cross-validation. The hyperparameters were re-tuned on each iteration. The hyperparameters from the fold with the best performance were then used to fit a final model to the entire training set. To assess predictive power in novel conditions, the model was also assessed using data from G1, G2 & S phase conditions. Note that for the acetylation predictions during the cell cycle, no additional training data was available for the G0 phase (in contrast to phosphorylation)

To assess the impact of using other ML algorithms on CAROM accuracy, additional models were built using Random Forests and AdaBoost. Similar accuracy to XGBoost was obtained using these approaches (S. Figure 22) [79]. AdaBoost is also a gradient boosting algorithm that can use decision trees as its base learners. For each learner, weights are assigned to its errors and these weights are used to adjust the next learner’s predictions.

For model interpretation, a single decision tree model was created to visualize the typical prediction path that an observation follows when its class is being decided. The decision tree was built using the scikit-learn Python package. The decision tree was trained on the entire dataset and the RandomizedSearchCV function was used to tune hyperparameters, including maximum depth. To address the class imbalance, synthetic minority oversampling (SMOTE) was used for training the decision tree model.

To build the ML model, each gene-reaction pair is assigned a class of −1, 0, or 1, corresponding to phosphorylated, unregulated and acetylated, respectively. For cases where genes/proteins were regulated by both PTM types in the training data, phosphorylation was assigned, as this was the minority class. This overlap occurred in 25 gene-reaction pairs in the *E. coli* dataset, 67 pairs for yeast and 2 for HeLa. Any genes that were included in the metabolic network, but not found in the corresponding PTM dataset, were assumed to be non-regulated. Any missing feature data was replaced with the median value. To account for the differences between organism characteristics, we normalized the features for each condition table on a scale of 0 to 1 for each condition. The catalytic activity and PFBA flux features showed unique organism-specific signatures when normalized, so these two attributes were scaled using their mean values. Reaction reversibility is a binary variable and therefore was not scaled. Prior to scaling, the maximum and minimum reaction flux features were limited to 100 to reduce feature range, as opposed to the value of 900 used in the statistical portion of the study. This step did not significantly affect the model accuracy (S. Figure 23)

Proteins that were not annotated to be acetylated or phosphorylated in any condition in the protein lysine modification database or the UniProt database were removed from the ML model [80,81]. However, this step did not significantly alter the accuracy as most metabolic proteins were annotated to be regulated by these PTMs (S. Figure 24). The final data used to train the CAROM-ML model included 2427 gene-reaction pairs for *E. coli*, 3039 for yeast, 3661 for HeLa, and 3582 for the G0 condition of the mammalian cell cycle dataset, for a total of 12,709 observations (S. Figure 25, S. Tables 15-17). The validation set, which includes the G1, S, and G2 phases, contained 10746 pairs (3582 for each phase).

### Shapley analysis

For determining features that have the largest influence in the ML models, we used the SHAP (SHapley Additive exPlanation) package in Python. SHAP uses the game theory concept of Shapley values for calculating each feature’s contribution to the model output [62]. The Shapley analysis was completed using TreeExplainer from the SHAP package. TreeExplainer is specifically designed for use with tree-based models. The Shapley value represents the average impact for each feature across for all possible orderings. This process is represented by the following equation:

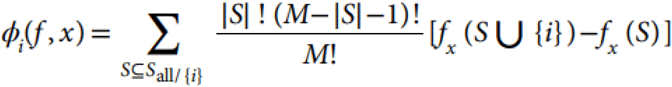

The Shapley value is the *ϕ*_*i*_(*f*, *x*) term, or the effect that feature *i* has on model *f*, given the independent variable data, *x*. *M* is the total number of features, and *M!* represents the number of possible feature combinations. *S* is a subset of the features excluding feature *i, |S|* is the number of features in subset *S*, and *f_x_(S)* is the model output for subset *S*. The SHAP values are relative to the average model output, called the base value. The base value can also be thought of as the null model output. Therefore, the sum of the SHAP values for a given observation is equal to the difference between the model prediction and the base value. Considering the SHAP values across all observations in a dataset provides insight into the overall feature importance, direction of a feature’s impact on the model output and relationships between the predictor features. For model interpretation using SHAP, the final XGBoost model and its training data were used as inputs to the TreeExplainer function.

## Author contributions

S.C conceived the study, S.C, K.S, H.L and F.S designed and performed research, and S.C wrote the manuscript with inputs from K.S and H.L.

## Data and materials availability

All datasets are available in the supplementary materials.

