## Supplementary Figures for "Metabolic signatures of regulation by phosphorylation and acetylation"

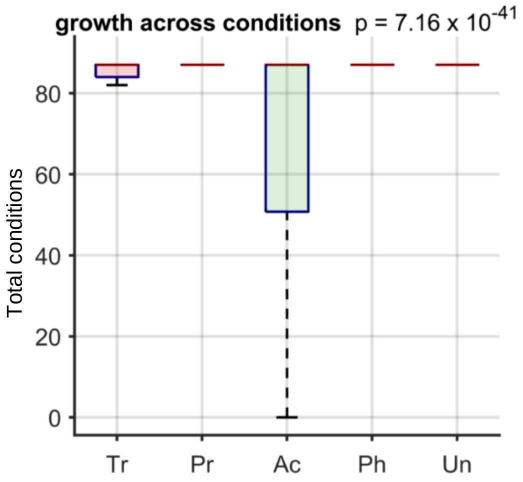


**S. Figure 1. Distribution of regulation based on gene essentiality across 87 different conditions.** These conditions comprise 56 different carbon sources including glucose, and 31 different nitrogen sources including ammonium ions. The total number of conditions in which each gene deletion was viable was calculated. This total number was then compared between targets of each regulatory mechanism. The box plots show that acetylation preferentially regulates the genes that impact growth across the 87 conditions. The box plot whiskers extend to the 99.3rd percentile of each distribution. The ANOVA p-value comparing the means is $7.1 \times 10^{-41}$.

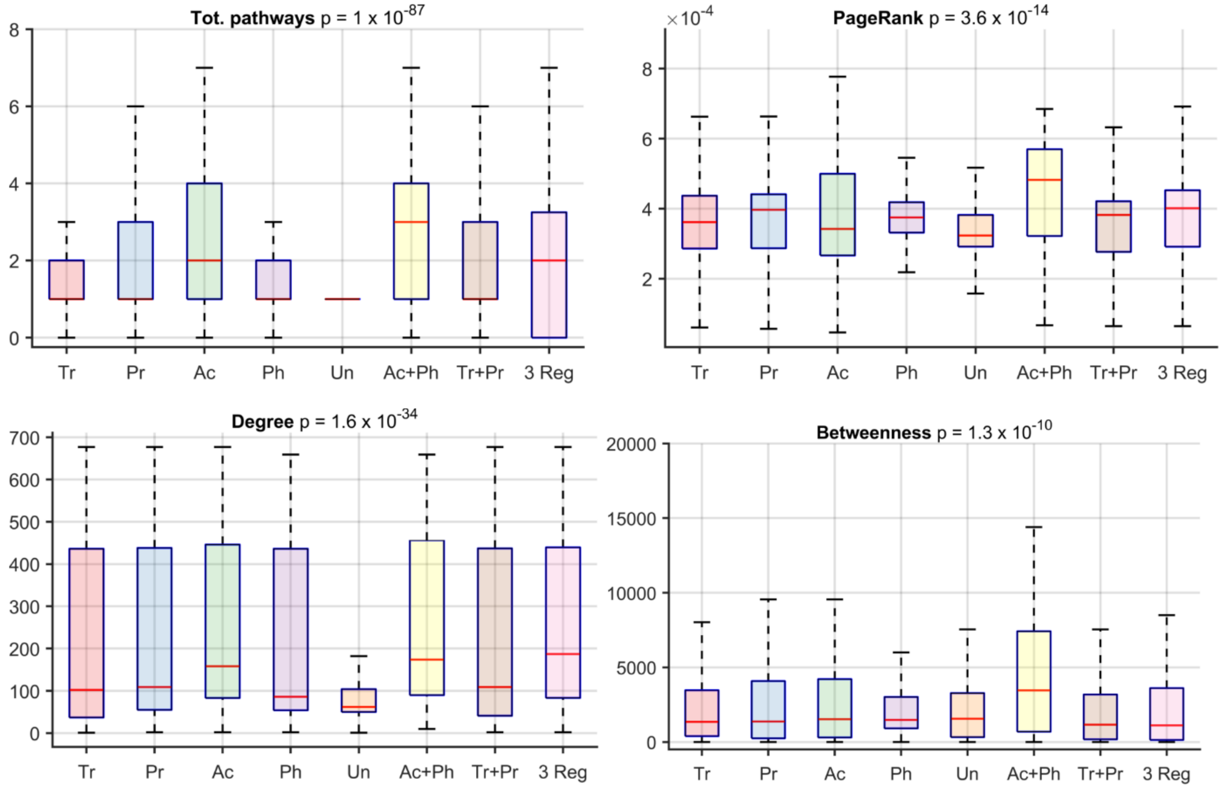


**S. Figure 2**. **Distribution of regulation based on topological properties of each reaction.** Four different topological properties are shown in the box plots - the total number of annotated pathways each reaction participates (Tot. pathways), the number of times each reaction is traversed during a random walk between reactions in the network (Pagerank), the total number of connected reactions (Degree) and the number of times each reaction appears on a shortest path between two reactions (Betweenness). These show that reactions that are regulated by any mechanism have a higher connectivity compared to those that are unregulated. Furthermore, reactions regulated by both acetylation and phosphorylation had the highest connectivity across all metrics. The ANOVA p-value comparing the means is provided in the title. (Abbreviations: regulation by both transcription and post-transcription (Tr + Pr), both acetylation and phosphorylation (Ac + Ph), at least 3 regulators (3 Reg), and Unregulated (Un)).

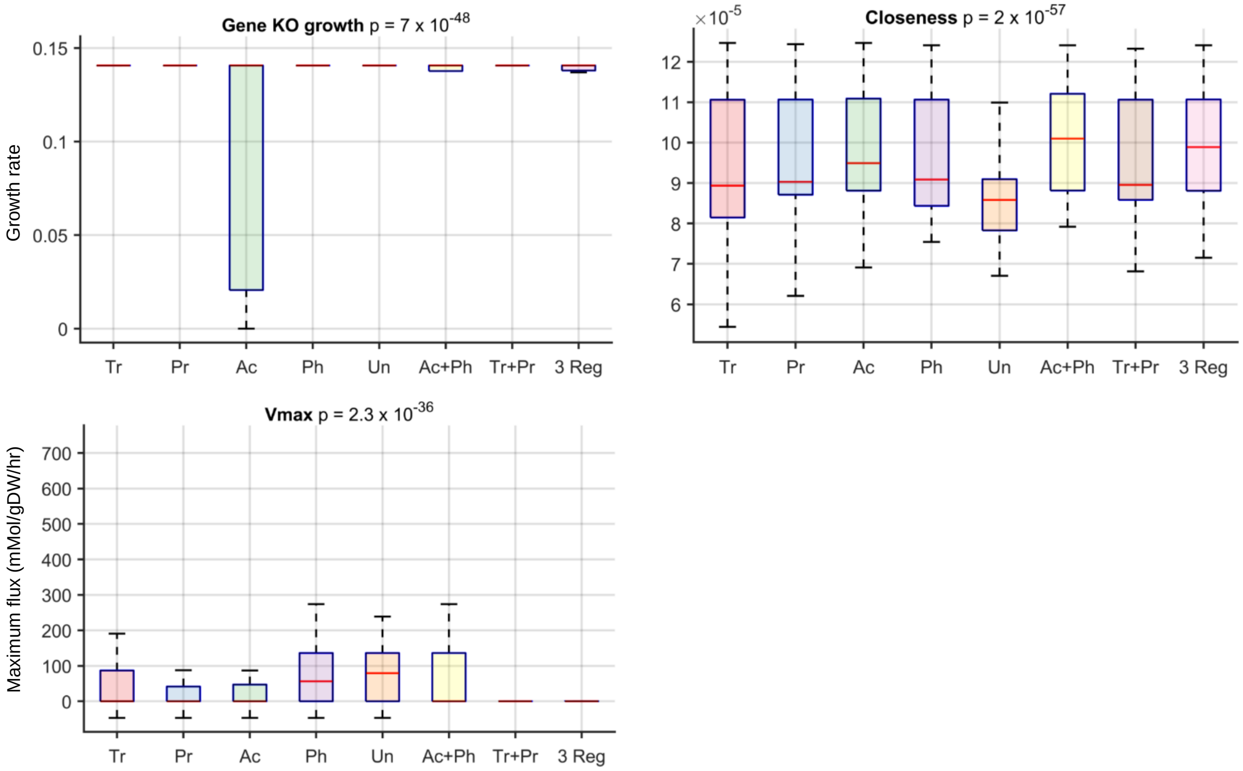


**S. Figure 3. Properties of reactions regulated by multiple mechanisms.** The box plots compare the properties of enzymes regulated by transcription, post-transcription, acetylation, phosphorylation with those regulated by both transcription and post-transcription (Tr + Pr), both acetylation and phosphorylation (Ac + Ph), or at least 3 regulators (3 Reg). This set of combinations among regulators was chosen as both acetylation and phosphorylation are PTMs, and the transcriptome and proteome of yeast cells show significant correlation. Reactions regulated by both acetylation and phosphorylation had the highest connectivity as measured by the inverse sum of the distance from a reaction to all other reactions in the network (Closeness). Apart from connectivity, reactions regulated by two different mechanisms did not share properties of reactions regulated by each individual mechanism. For example, reactions regulated by acetylation and phosphorylation were not likely to be essential or have high maximum flux. The ANOVA p-value comparing the means is provided in the title.

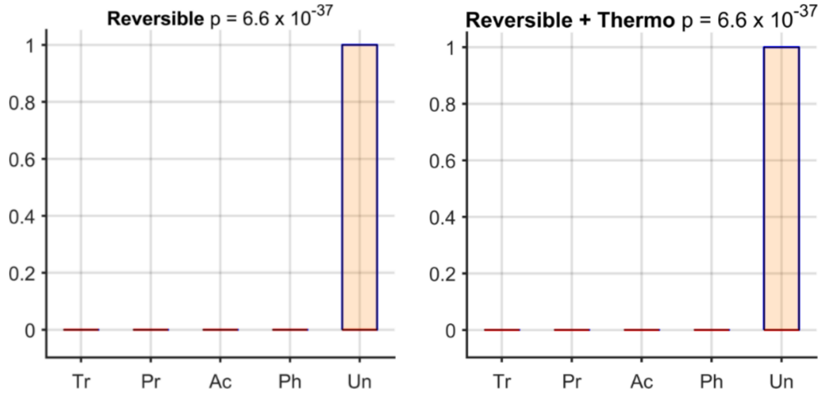


**S. Figure 4. Distribution of regulation based on reaction reversibility.** Reversible reactions were highly likely to be not regulated by any of the four mechanisms. The left panel compares the distribution of regulation of reversible reactions based on the annotation from the Yeast 7 model (reversible reactions are set to 1 and irreversible reactions are set to 0). The panel on the right uses an updated list based on thermodynamic analysis of the Yeast metabolic model by Martinez *et al* [49].

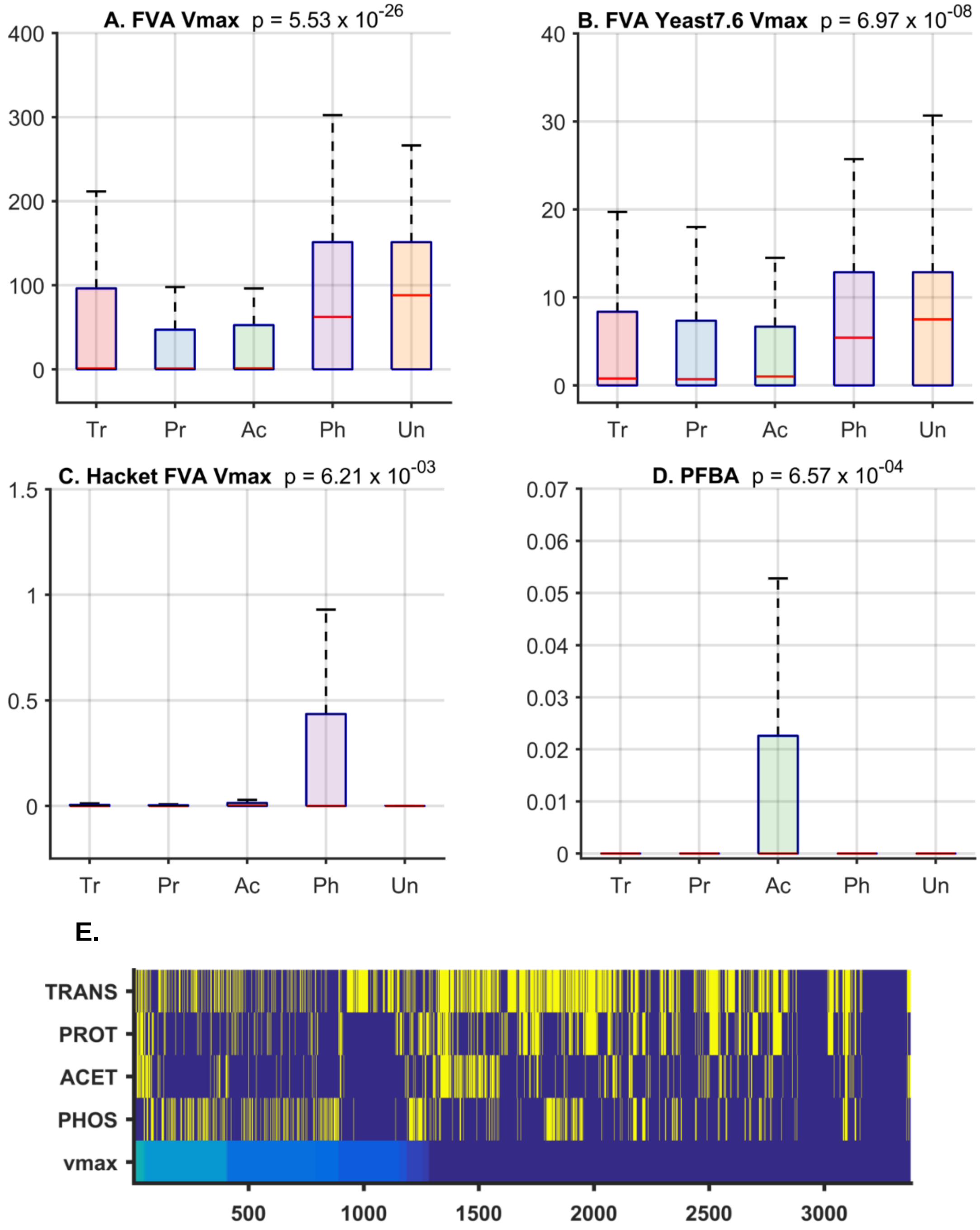


**S. Figure 5. Distribution of regulation based on magnitude of maximum possible flux** (mmol/gDW/hr) through each reaction. The plots compare the distribution of regulation using flux calculated using various methods and models. The ANOVA p-value comparing the means is provided in the panel title of each plot. These results show that phosphorylated reactions are highly enriched among those reactions with high maximum flux. **A.** Maximum flux through each reaction was calculated using FVA using the Yeast 7 model without assuming that cells maximize their biomass (the default objective in FVA and FBA). The box plots compare the maximum flux value of reactions regulated by each mechanism. **B.** Maximum flux through each reaction was calculated using FVA without assuming that cells maximize their biomass using the Yeast 7.6 model (Yeast 7 model was used for all analyses).

**C.** The flux through the model was first fit to the experimentally inferred flux data from Hackett *et al*[21]. The maximum flux through all reactions was then determined using FVA. **D.** The flux through each reaction was inferred from Parsimonious FBA (PFBA). Note that PFBA does not provide the maximum flux but the flux value that minimizes the sum of flux through all reactions while maximizing the biomass objective. Hence it does not reveal any futile cycles or redundancy in the network. **E.** The heatmap shows the distribution of regulation based on magnitude of maximum possible flux (Vmax) through of each reaction. Reactions are sorted based on Vmax inferred from FVA. The columns correspond to each reaction-gene pair. Those that are regulated by each mechanism are shown in yellow, while those that are not regulated by a specific mechanism are in blue.

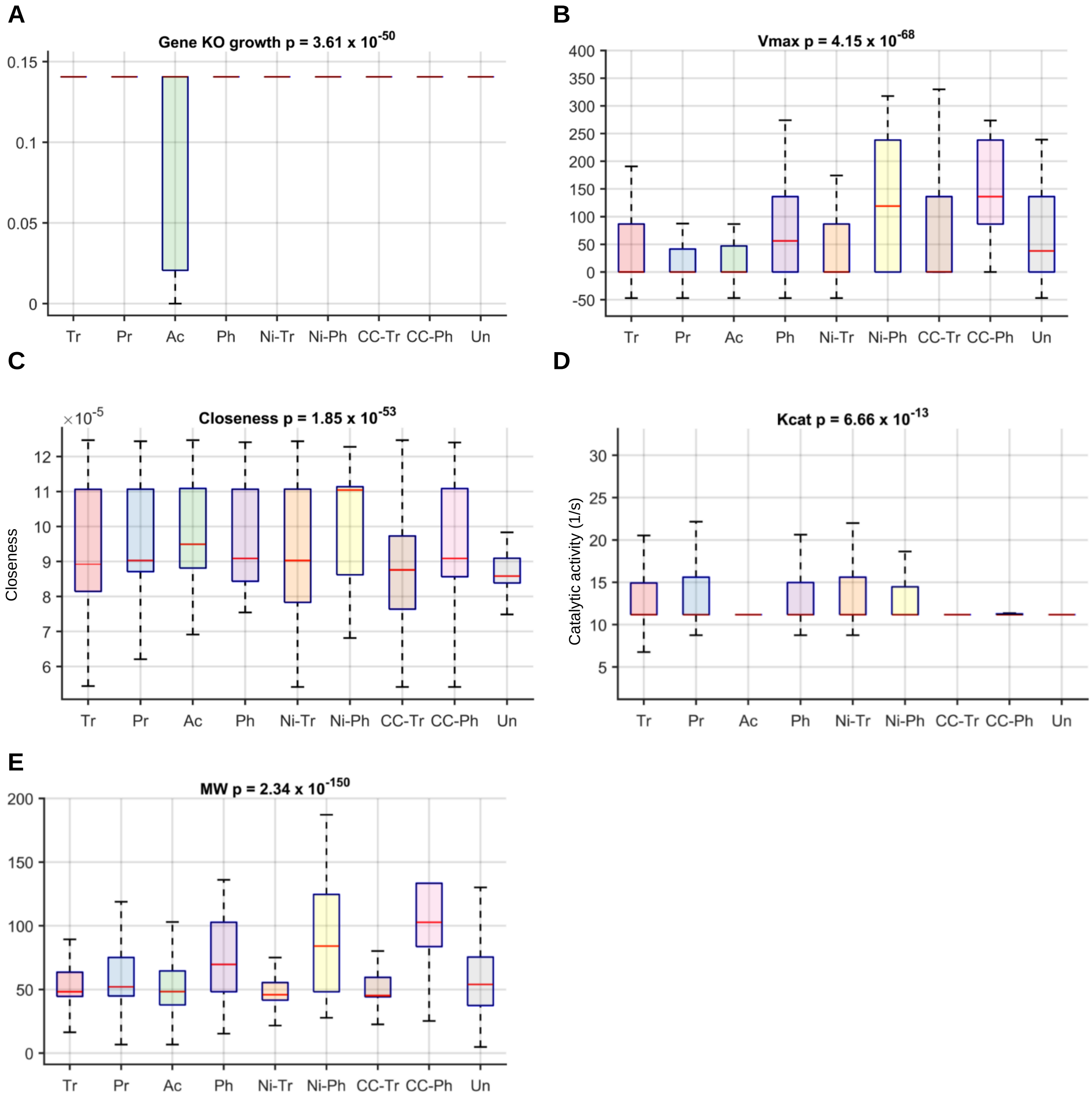


**S. Figure 6. Comparison of the properties of enzymes in yeast regulated by each mechanism during the cell cycle (CC-Tr, CC-Ph) and nitrogen starvation (Ni-Tr, Ni-Ph)**. Data from stationary phase conditions (transcription (Tr), post-transcription (Pr), acetylation (Ac), phosphorylation (Ph) or Unregulated (Un)) are shown for comparison. Similar to stationary phase, enzymes that impact growth when knocked out are likely to be acetylated **(A),** enzymes that catalyze reactions with high flux are likely to be regulated through phosphorylation in all three conditions **(B),** enzymes that are highly connected are likely to be regulated by one of the four mechanisms **(C).** No consistent difference across datasets was observed in regulation based on the enzyme catalytic activity (kcat) of the target enzyme **(D)** and enzymes regulated by phosphorylation on average tend to have high molecular weight **(E)**. The Anova p-value comparing the differences in means is shown in the title.

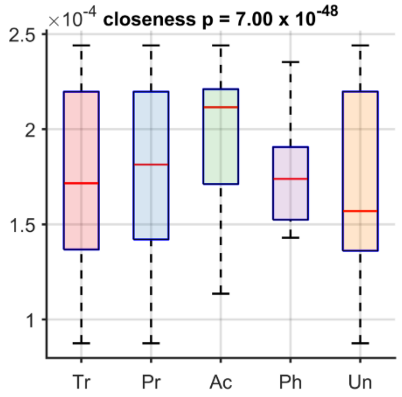


**S. Figure 7. Comparison of the topological properties of enzymes in *E. coli* regulated by each mechanism.** The Anova p-value comparing the differences in means is shown in the title. Similar to yeast, enzymes that are highly connected (i.e. high closeness) are likely to be regulated.

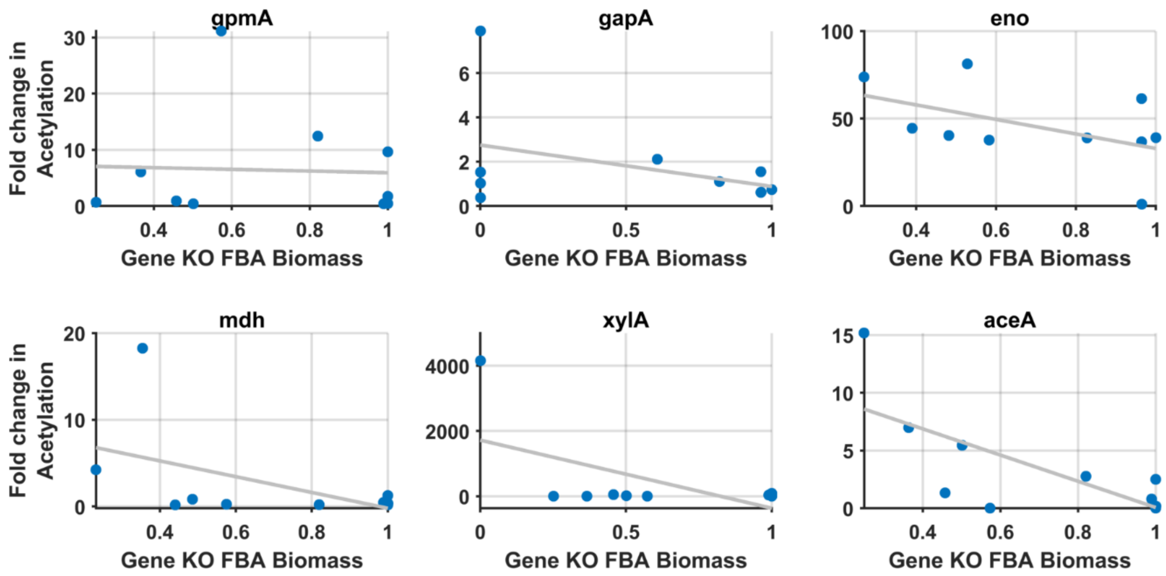


**S. Figure 8. Condition-specific essentiality is correlated with acetylation.** The scatter plots show the association between the impact of a gene knockout on biomass from FBA with the acetylation levels of the corresponding protein in a given condition. On average, increased essentiality is associated with an increase in acetylation. All proteins with at least 2 fold change in acetylation between conditions and are part of the metabolic model are shown. The change in biomass relative to glucose is show in the x-axis. The correlations were observed even when the total absolute acetylation levels were considered instead of relative levels to proteins.

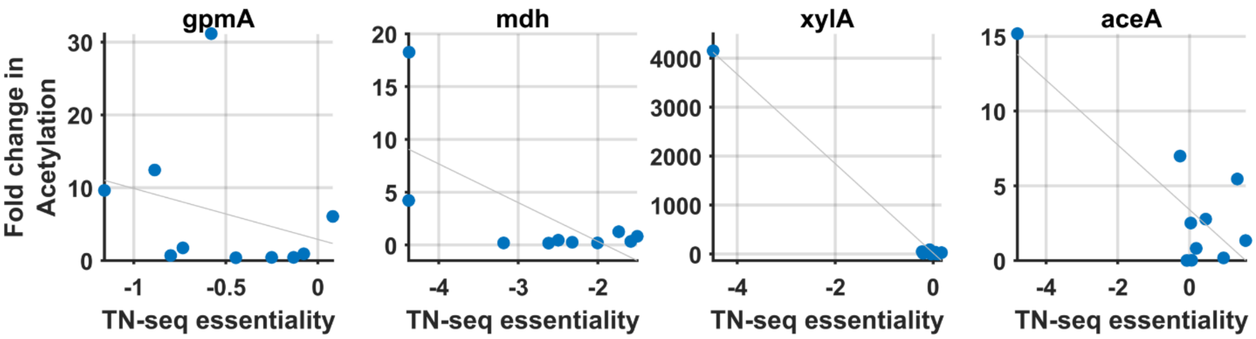


**S. Figure 9. Condition-specific essentiality from TN-seq is correlated with acetylation.** The scatter plots show the association between the impact of a gene knockout on viability from Transposon mutagenesis screens with the acetylation levels of the corresponding protein in a given condition. All proteins in the metabolic model with available TN-seq data and acetylation data across conditions from Schmidt et al study are shown. Although FBA made false positive growth predictions for some enzymes such as XylA (S. Figure 8), our results were observed even with experimentally derived knockout screens, suggesting that this link between essentiality and acetylation is robust.

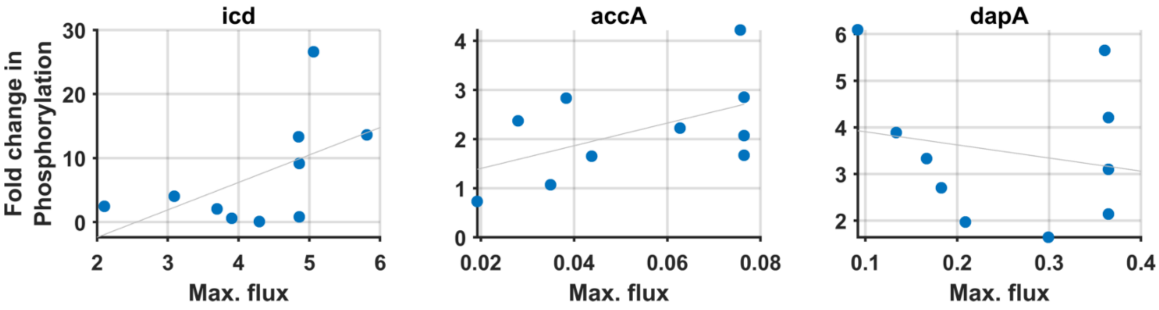


**S. Figure 10. Correlation between maximum flux and phosphorylation levels** (normalized to glucose). All proteins that showed at least 2-fold change in phosphorylation levels between conditions are shown. This trend was observed with both the total phosphorylation levels and relative levels normalized to proteins. While in most cases a change in maximal flux or essentiality resulted in a change in regulation by PTMs (Figure 2F), there were exceptions. For example, dapA did not show this trend suggesting that other factors likely influence regulation by PTMs in a combinatorial fashion.

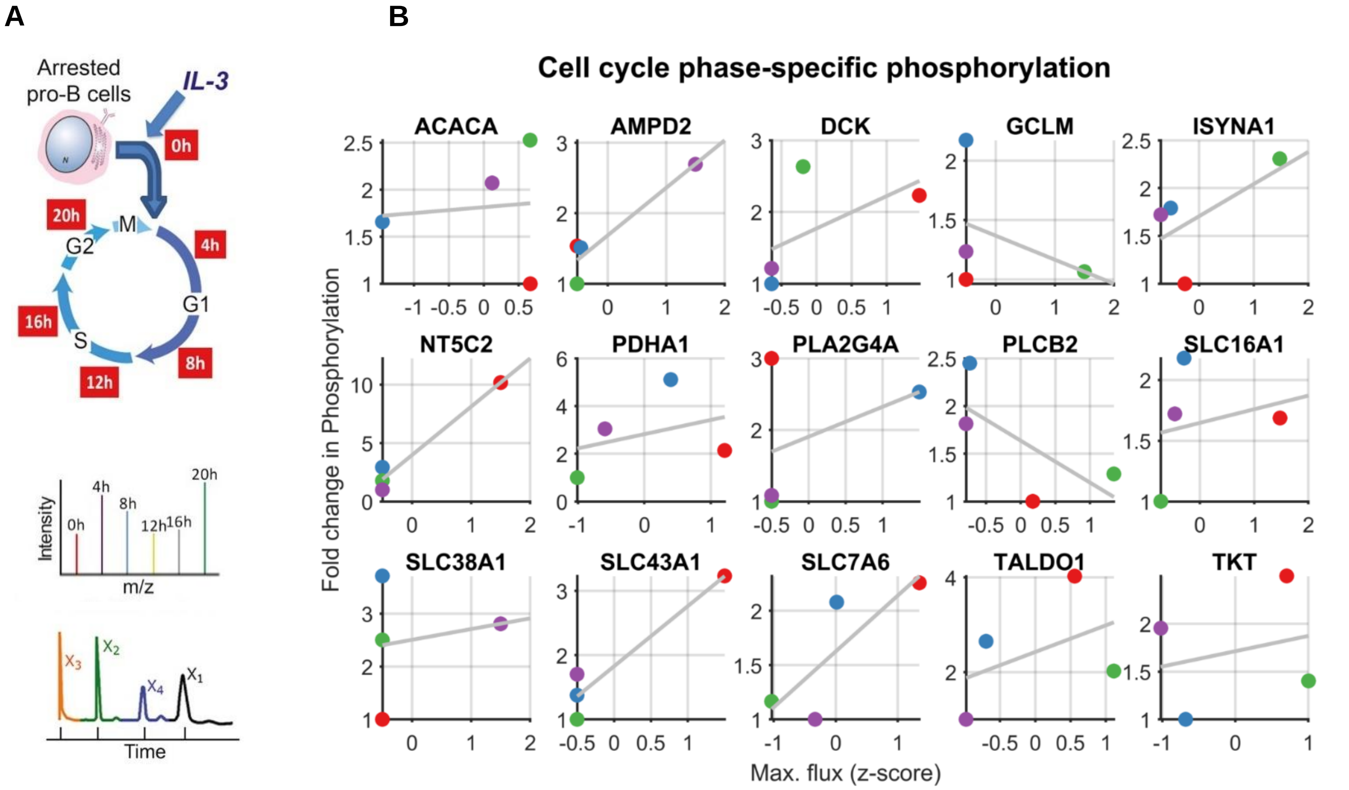


**S. Figure 11. Enzyme phosphorylation across the cell cycle correlates with maximum flux**. **A.** Pro-B cells were synchronized by growing in media without IL-3 for 36 hr and then released from G0 by re-growing them in the presence of IL-3. Phospho-proteomics data were collected at each phase. In addition, metabolomics data from the same system from Lee-et al was used to build metabolic models for each phase. **B**. All proteins that show at least two-fold change in phosphorylation are shown. For those proteins associated with multiple reactions, the reaction with the highest flux change is shown. Markers are colored by cell cycle phase (red G0, blue G1, green S, violet G2). Among the proteins that showed at least 5-fold difference in phosphorylation between phases, the average correlation was 0.56 between the normalized maximum flux and fold change in phosphorylation. The correlation increases to 0.9 for proteins that show a tenfold difference and is 0.17 for proteins that show 2-fold difference.

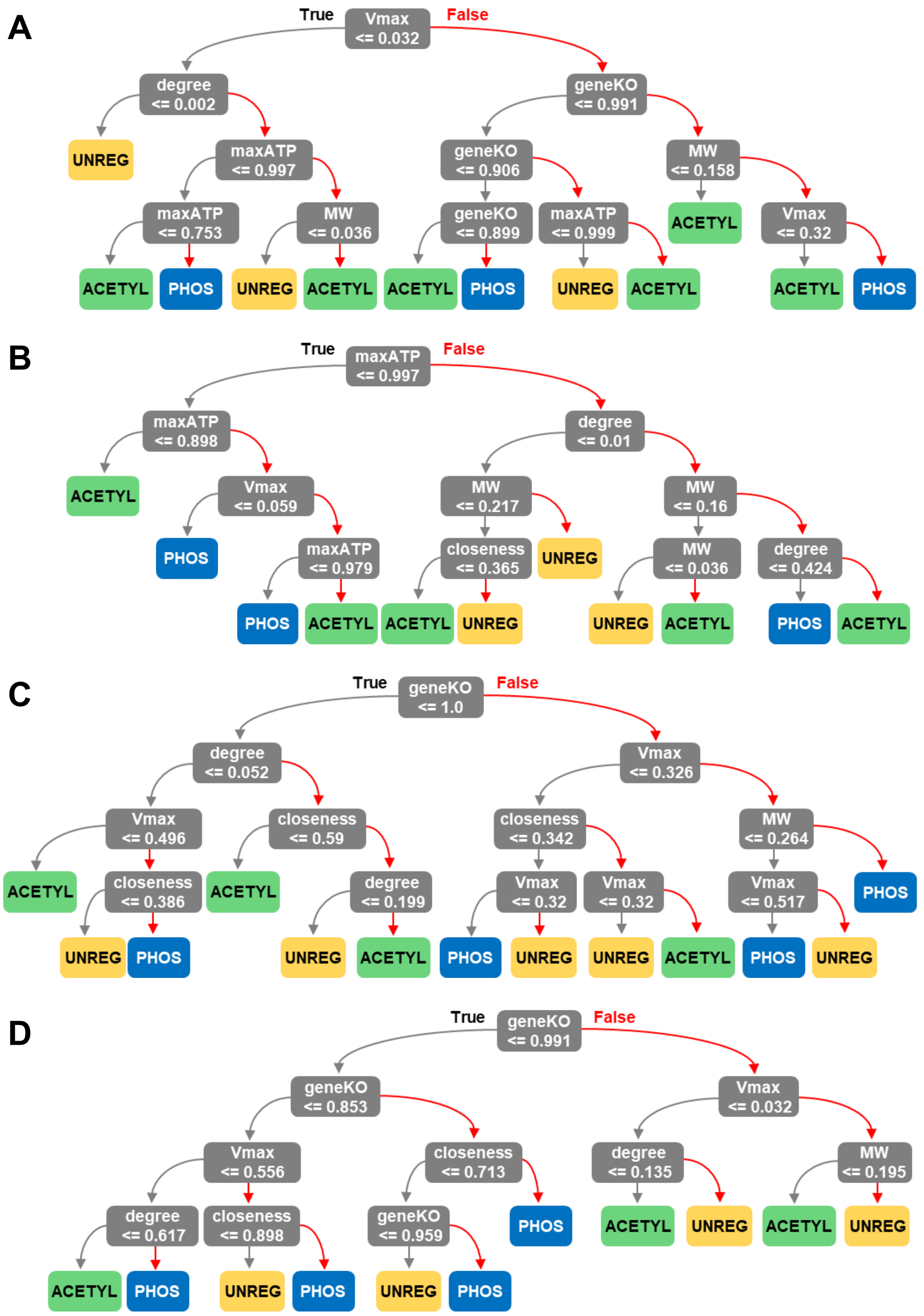


**S. Figure 12. Representative decision trees with maximum depth of 4**. Single decision tree models were trained for the multi-organism **(A)**, *E. coli* **(B)**, yeast **(C)**, and mammalian **(D)** datasets. Only the top 50% most important features, as identified in the Shapley analysis, were used to train the trees.

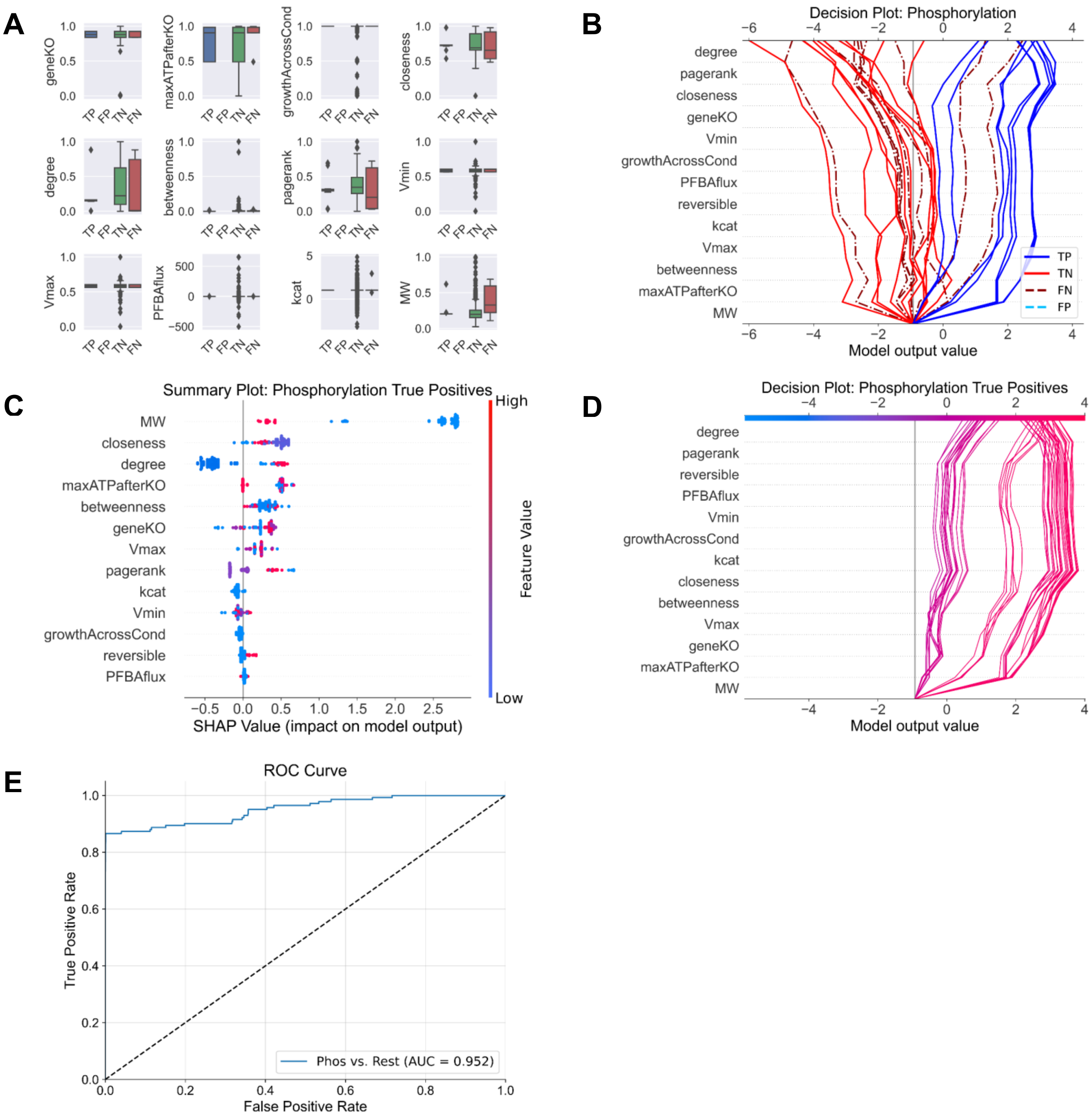


**S. Figure 13. Analysis of model predictions on the cell-cycle phosphorylation data** (refer to Figure 4 in the main text). **A.** Feature distributions for phosphorylated gene-reaction pairs are compared between true positive (TP), true negative (TN) and false negative (FN) observations using boxplots. There were no false positives from this validation test. **B** SHAP decision plot was created for 50 random observations to compare trends between the classification groups. Values on the x-axis represent log odds of belonging to the phosphorylation class. **C** and **D.** The phosphorylated gene-reaction pairs that were correctly classified (true positives) are displayed in a SHAP summary plot (C) and decision plot (D). **E**. ROC curve for the model's phosphorylation predictions on the cell-cycle data.

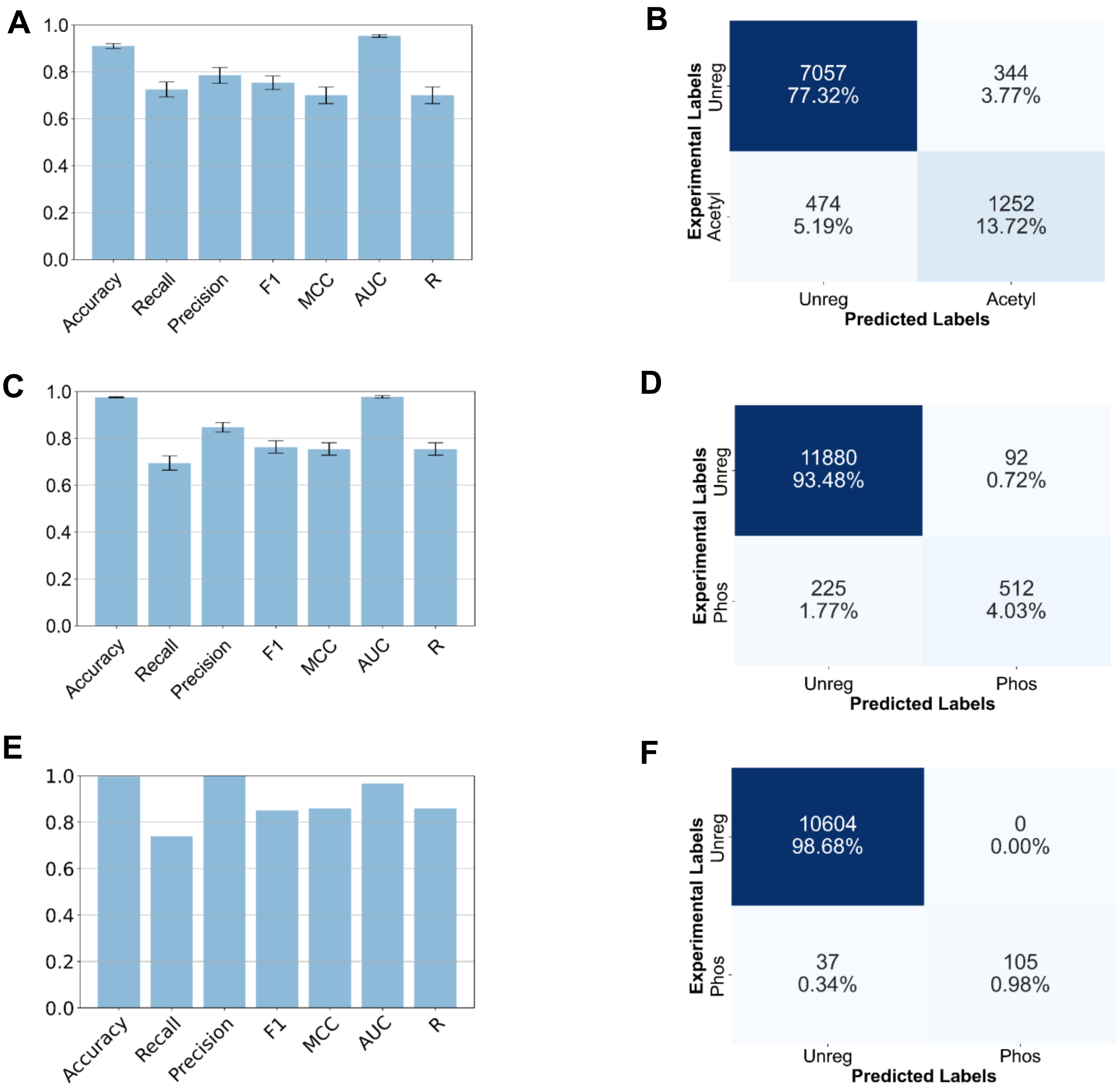


**S. Figure 14. Binary classification models for predicting acetylation and phosphorylation separately**. The pipeline for training the models was identical to process used for the multi-class model. **A, B** The 5-fold cross-validation results for the acetylation model. **C, D.** The 5-fold cross-validation results for the phosphorylation model. **E, F.** The phosphorylation model was used to predict the cell-cycle validation dataset, which includes the G1, S and G2 phases. Overall, these results show that the ternary classification model outperforms the binary classification models.

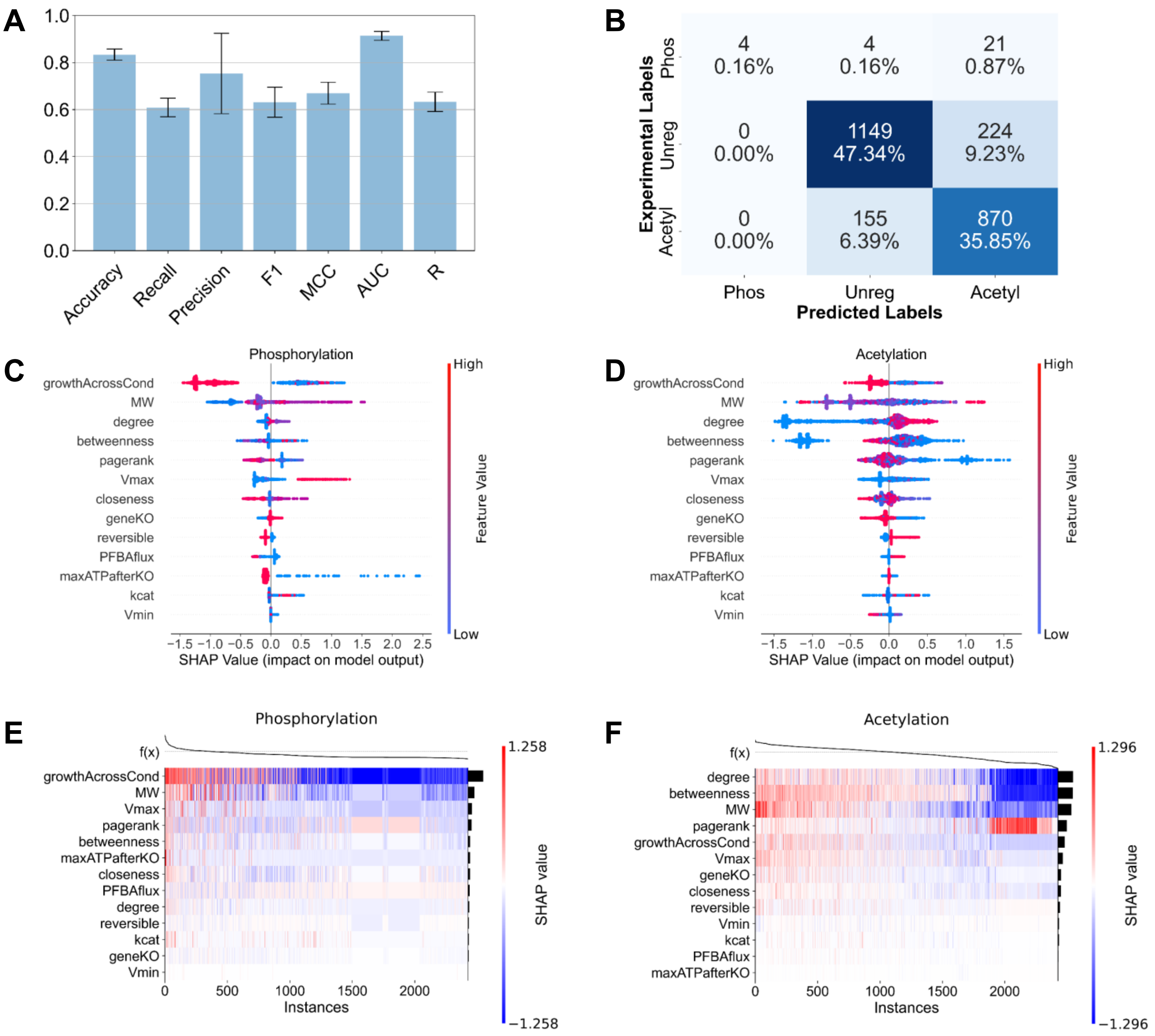


**S. Figure 15. Organism-specific ML models – E. coli.** XGBoost model trained on the *E. coli* dataset. **A, B.** 5-fold cross-validation results. **C, D.** SHAP value summary plots for the phosphorylation and acetylation classes. **E, F.** SHAP value heatmaps for the phosphorylation and acetylation classes. Observations are clustered by the model output, f(x).

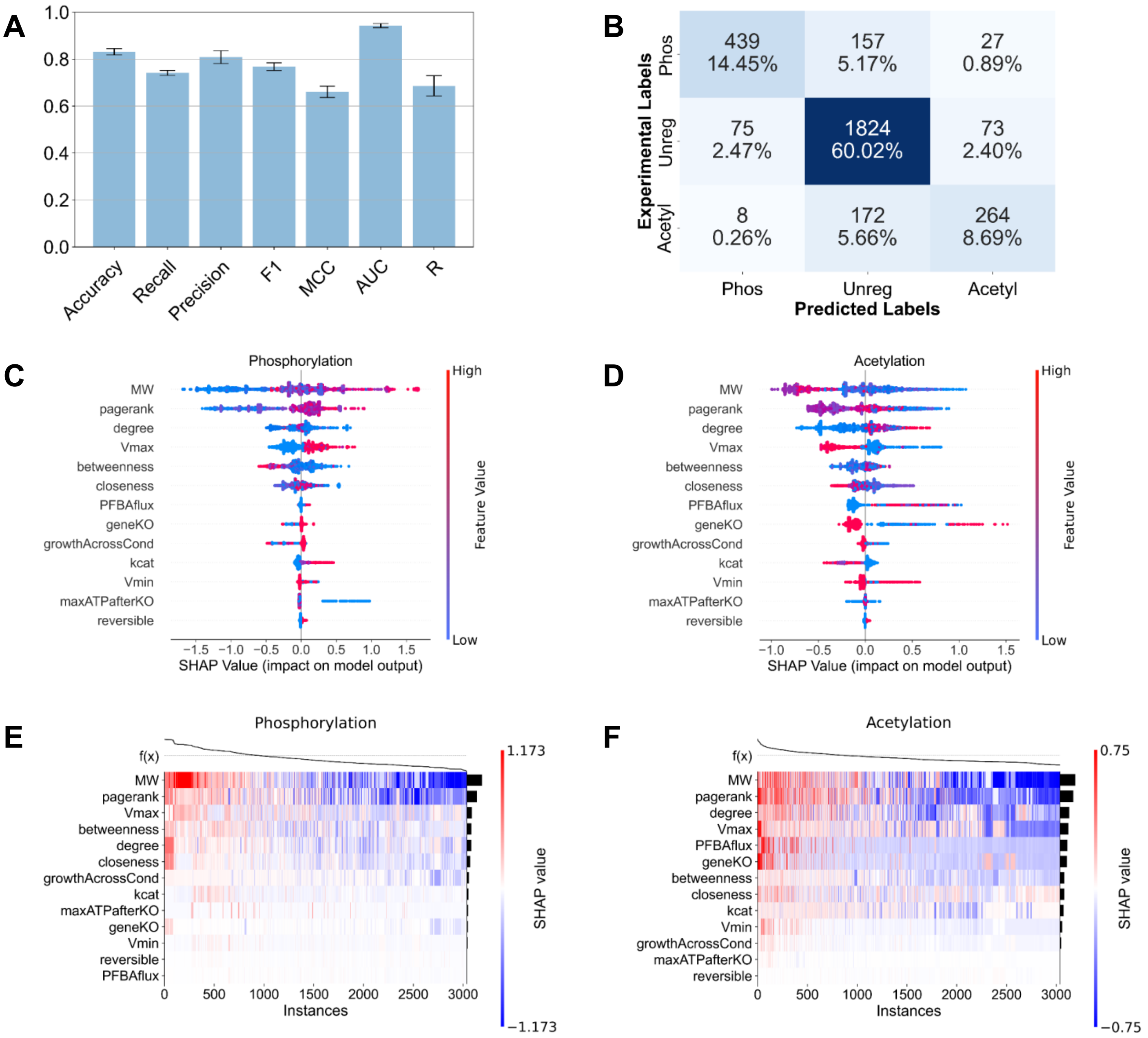


**S. Figure 16. Organism-specific ML models – S. cerevisiae.** XGBoost model trained on the yeast dataset. **A, B.** 5-fold cross-validation results. **C, D.** SHAP value summary plots for the phosphorylation and acetylation classes. **E, F.** SHAP value heatmaps for the phosphorylation and acetylation classes.

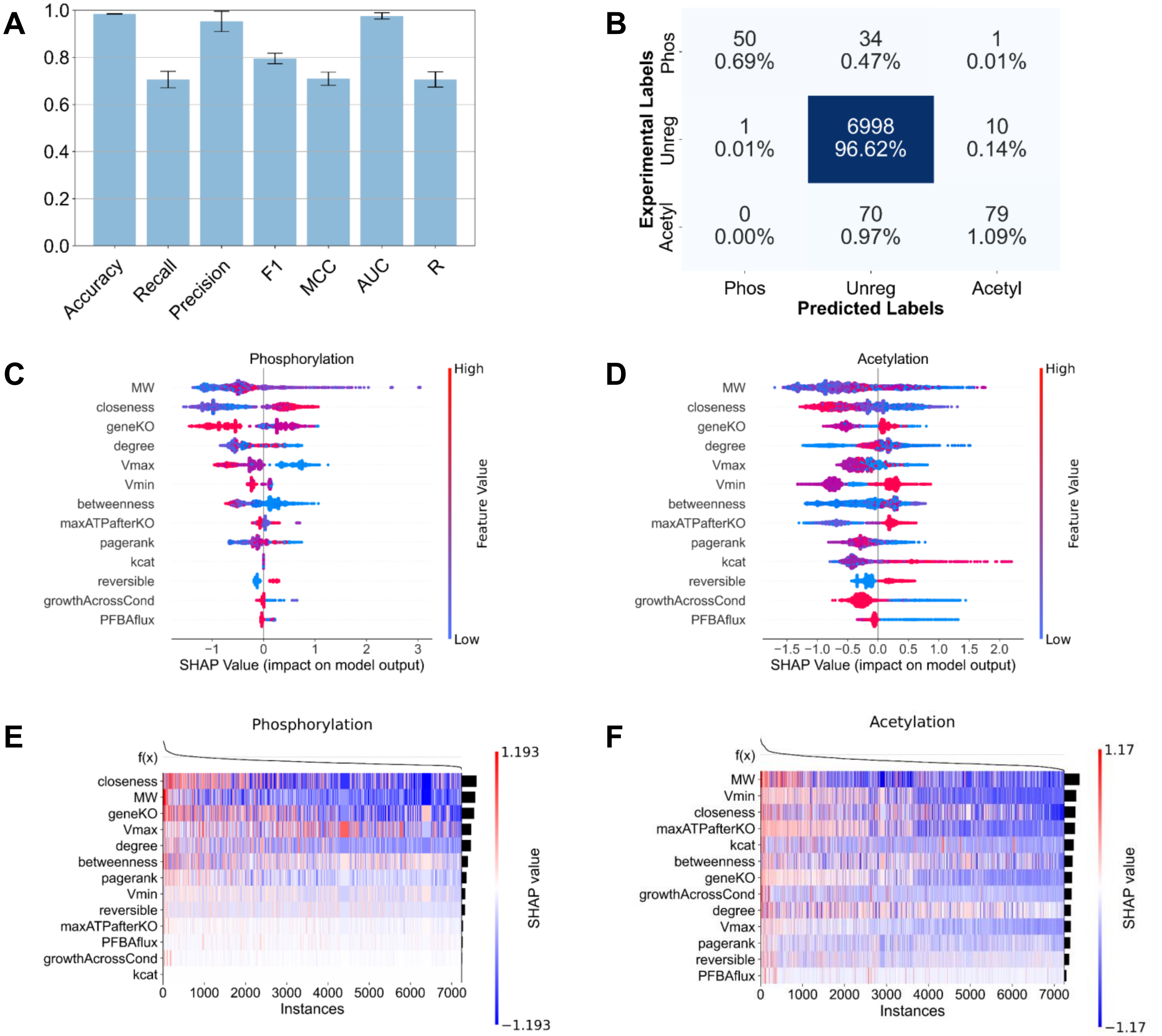


**S. Figure 17. Organism-specific ML models – mammalian cells.** XGBoost model trained on the mammalian dataset. **A, B.** 5-fold cross-validation results. **C, D.** SHAP value summary plots for the phosphorylation and acetylation classes. **E, F.** SHAP value heatmaps for the phosphorylation and acetylation classes.

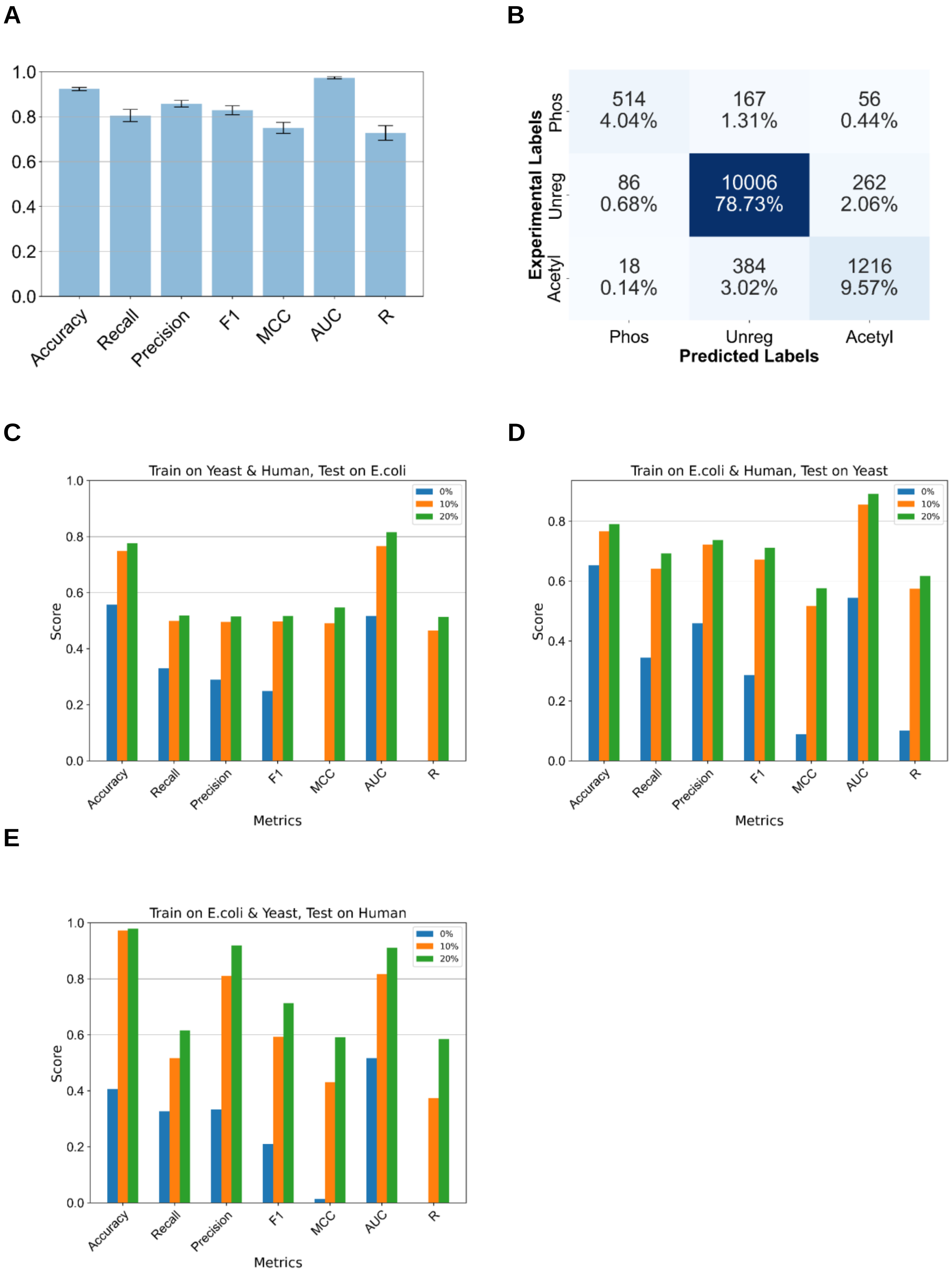
A
B
C
D
E
Accuracy
Recall
Precision
F1
MCC
AUC
R
Experimental Labels
Predicted Labels
Phos
Unreg
Acetyl
514
4.04%
167
1.31%
56
0.44%
86
0.68%
10006
78.73%
262
2.06%
18
0.14%
384
3.02%
1216
9.57%
Train on Yeast & Human, Test on E.coli
Train on E.coli & Human, Test on Yeast
Train on E.coli & Yeast, Test on Human
Score
Metrics
0%
10%
20%

**S. Figure 18. Impact of including organism type in the ML model and prediction on unseen organism data. (A, B).** 5-fold cross-validation results for XGBoost model with organism-type included in the training data. The organism type was added as a categorical array where a 1 designated *E. coli*, 2 for yeast and 3 for human. The cross-validation results were extremely consistent with those from the primary model, suggesting that the model's decision-making is not influenced by organism type. **(C, D, E).** CAROM models were trained on the data from two organisms and used to make predictions on the third (e.g. train on E. coli and yeast, test on mammalian). Data from the test organism was moved to the training data in increments of 0%, 10% and 20%. Model performance improved significantly after including a small number of samples from the test organism in the training dataset.

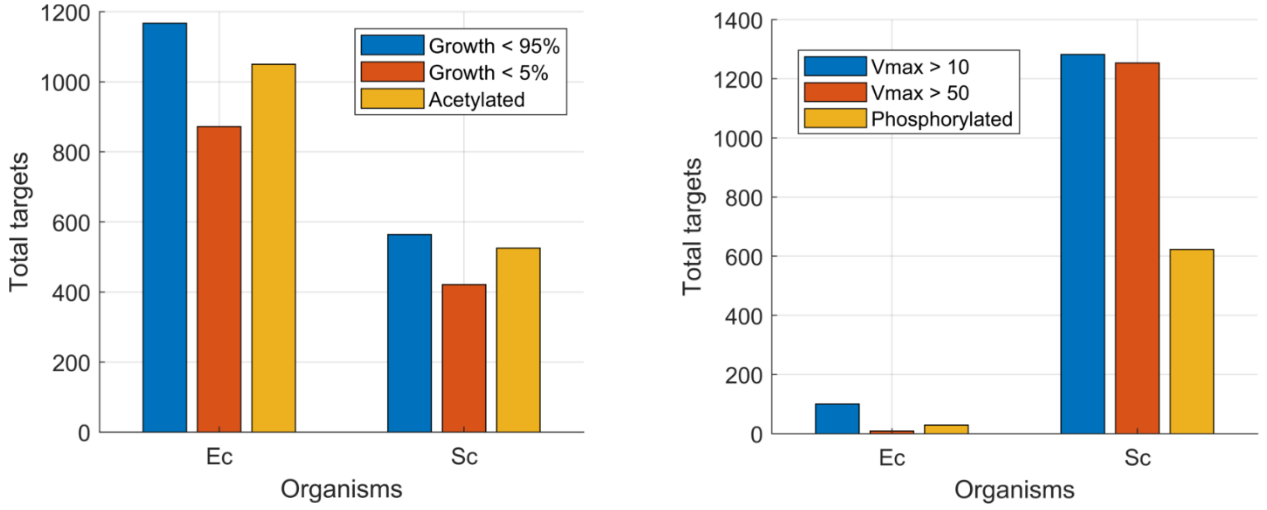


**S. Figure 19. Comparison of total number of targets between species.** Total number of regulation targets (i.e. gene-reactions) of PTMs in *E. coli* (Ec) and yeast (Sc) are compared with those that have high Vmax and are growth limiting in those species in the stationary phase condition.

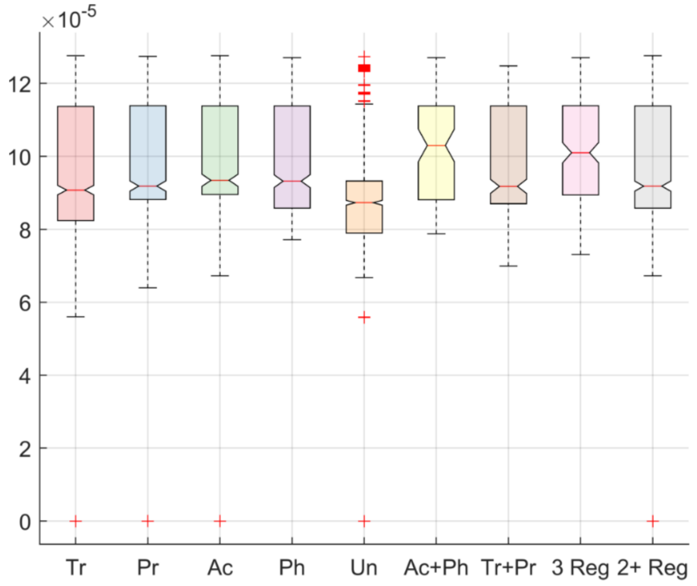


**S. Figure 20. Robustness of topological analysis.** Highly connected metabolites (ATP ADP AMP NADH NAD) were removed from the yeast model prior to the calculation of topological parameters. The box plots compare the properties of enzymes regulated by transcription (Tr), post-transcription (Pr), acetylation (Ac), phosphorylation (Ph), both transcription and post-transcription (Tr + Pr), both acetylation and phosphorylation (Ac + Ph), or at least 3 regulators (3 Reg). Reactions regulated by both acetylation and phosphorylation had the highest connectivity as measured by the Closeness. The ANOVA p-value comparing the means is 3e-46 for closeness, 2e-29 for degree (not shown) and 5e-15 for pagerank (not shown).

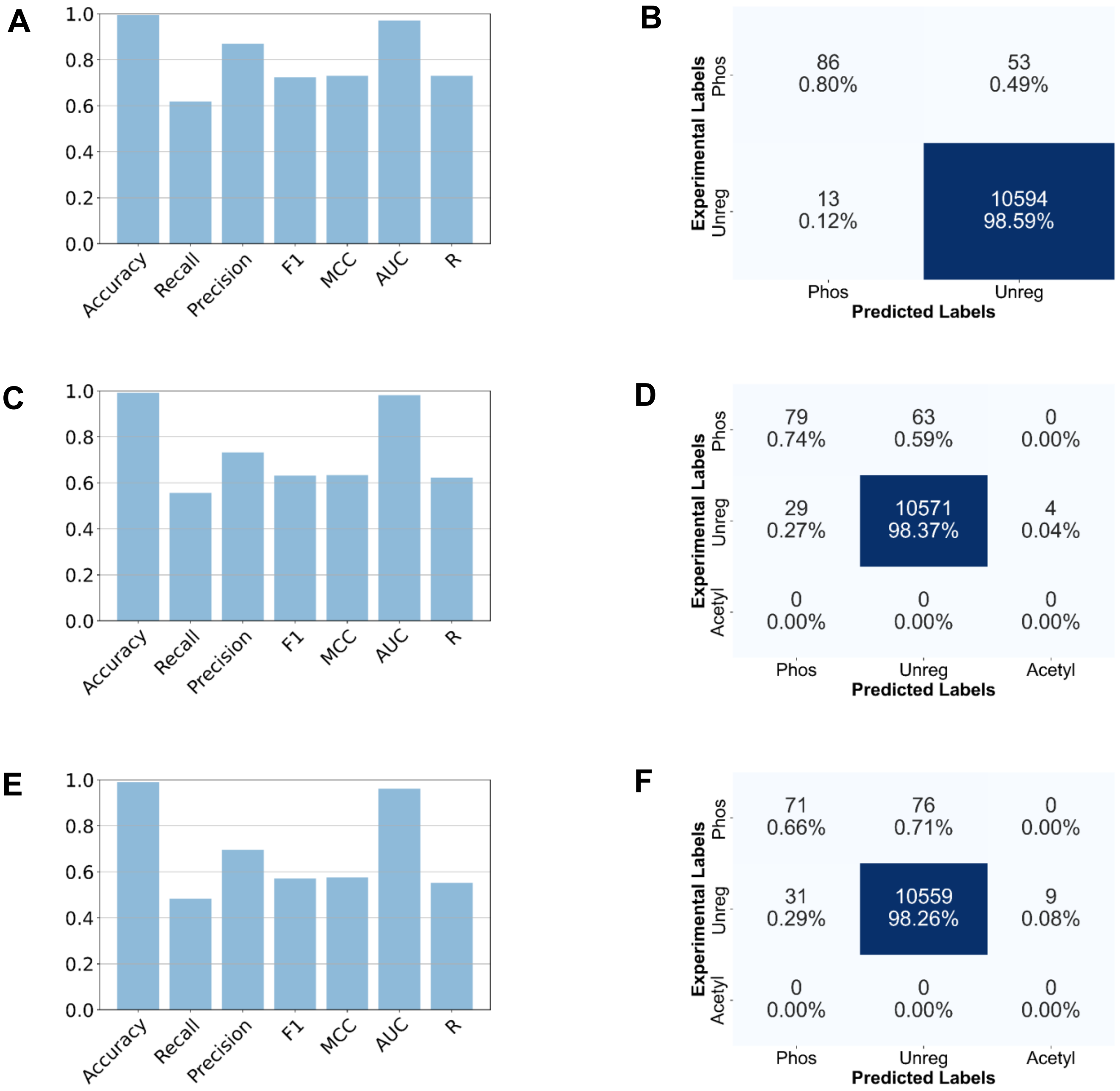


**S. Figure 21. Impact of training on different phases of the cell cycle.** Models were trained by replacing the G0 cell-cycle data from the training set with the feature matrix from the remaining phases: G1, S, and G2. Each model was then used to predict the phosphorylated genes from the phases not featured in the training. These results are shown here for the G1-model **(A, B)**, S-model **(C, D)** and G2-model **(E, F)**. All three models, especially for S and G2, performed inferior to the primary CAROM model in regard to this validation test. These results suggest that S and G2 conditions have a distinct phosphorylation pattern from the remaining conditions.

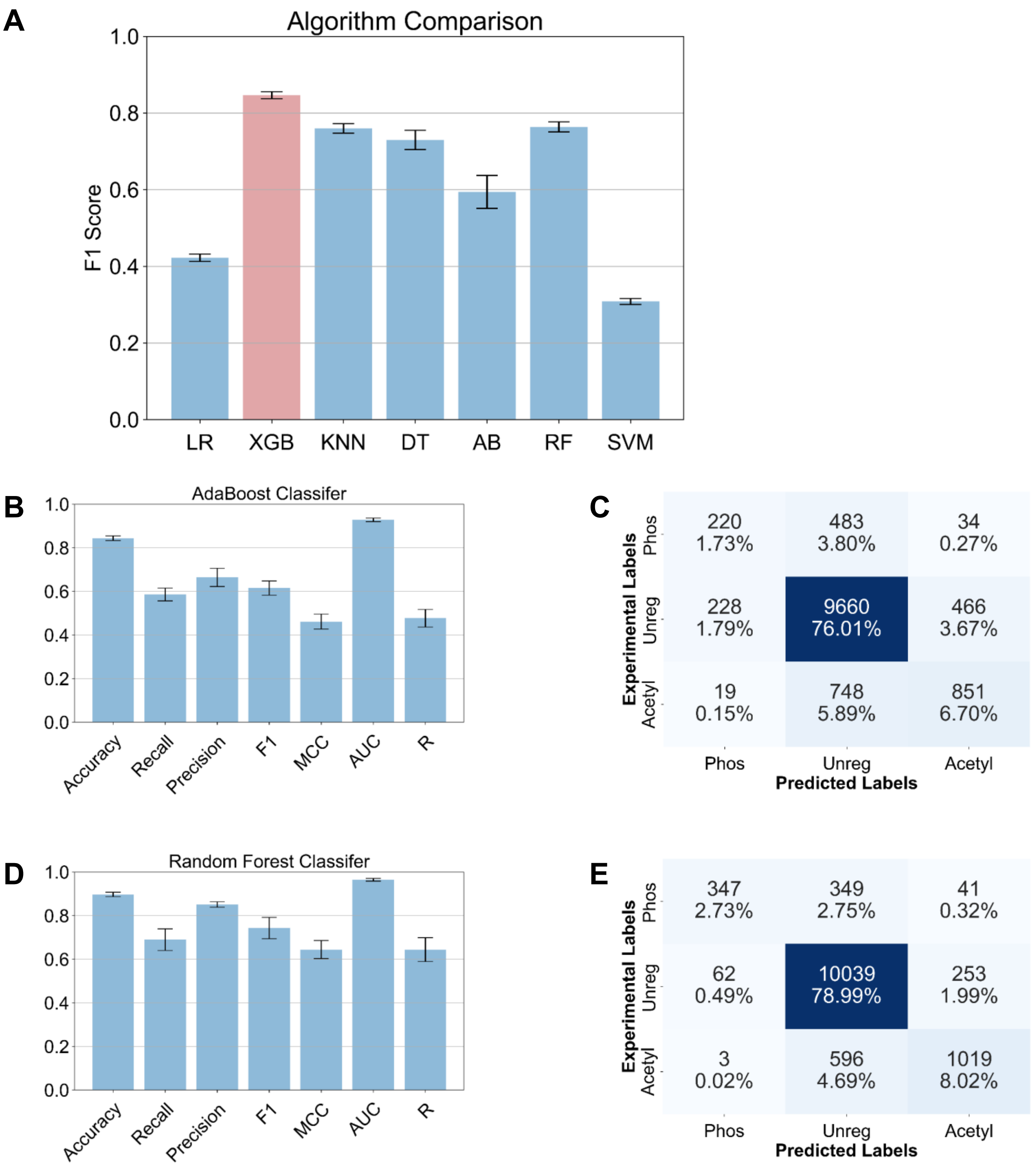


**S. Figure 22. CAROM model performance using various ML algorithms**. 5-fold cross-validation results were compared for various untuned algorithms, with F1 score used as the metric **(A)**. XGBoost, colored in red, had the best performance and was therefore used for the main CAROM model. AdaBoost **(B, C)** and random forest **(D, E)** models were further tested by tuning their hyperparameters and performing 5-fold cross-validation.

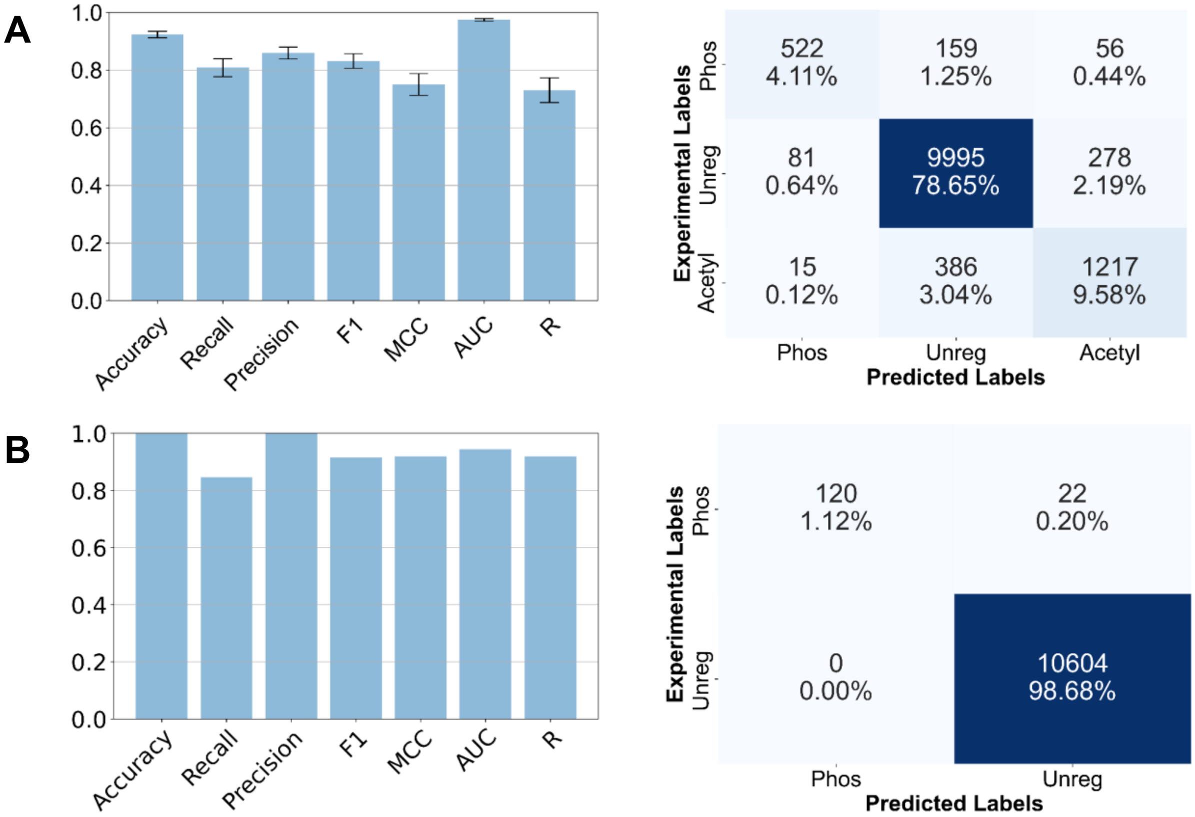


**S. Figure 23. Impact of increasing the maximum flux threshold on the ML results.** For the ML analysis, the Vmax and Vmin features were constrained to magnitudes below 100 in order to reduce the effect of unconstrained reactions and the variability across organism types. Here we show that the CAROM model is robust to increasing the threshold to the 900 mmol/gDW/hr value used for the ANOVA testing. A supplementary model was trained on the *E. coli*, yeast, HeLa and G0 phase data after adjusting this threshold. **A.** Results from training the model using 5-fold cross-validation. **B.** The model was used to predict on the cell cycle validation dataset, which includes the G1, S and G2 phases.

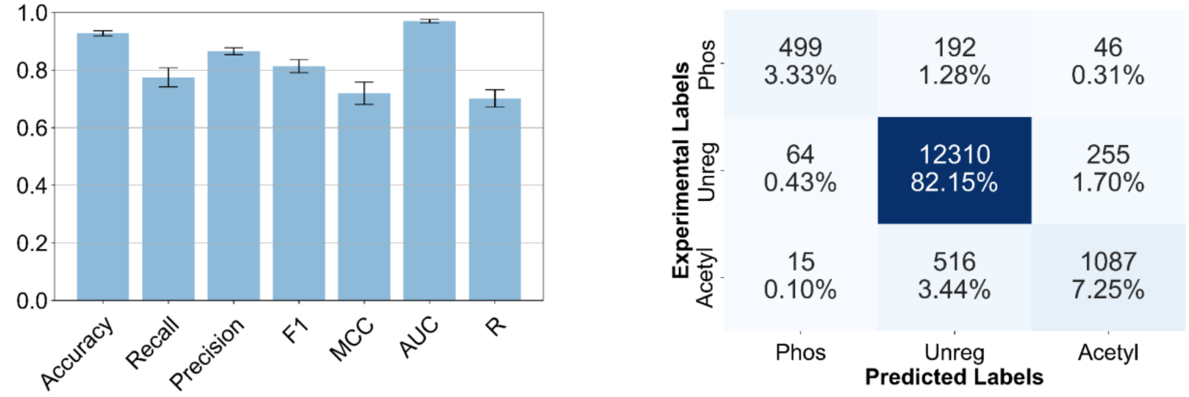


**S. Figure 24. Impact of retaining genes that do not have evidence for phosphorylation or acetylation.** 5-fold cross-validation results for model trained on full set of genes is shown. For the main CAROM model, online databases were used to compile a list of enzymes that have been found to be phosphorylated or acetylated in published studies. Non-annotated enzymes were removed from the training data. Here we show the results for the model which had these non-annotated enzymes included in the training data did not differ from the model with these genes removed during the model construction.

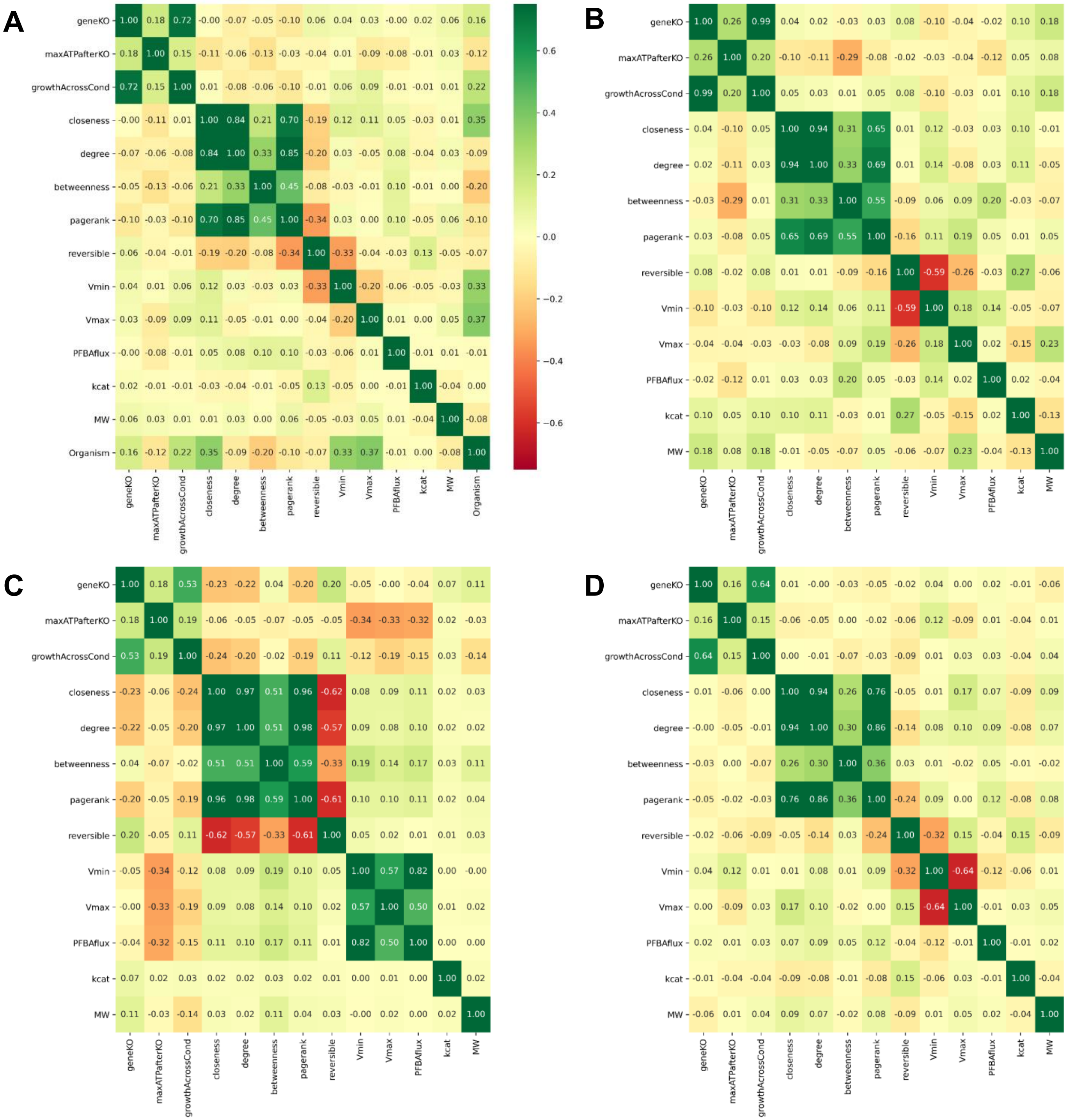


**S. Figure 25. Correlation map of all model features.** Heatmap of Pearson's correlation between feature values for the following datasets: all organism types **(A)**, yeast **(B)**, *E. coli* **(C)**, and human **(D)**.

### Supplementary Tables

**A**

| Regulatory mechanisms | | Reaction Overlap | p-value |
|---|---|---|---|
| TRANS | PROT | 421 | $4.12 \times 10^{-30}$ |
| TRANS | ACET | 285 | $2.36 \times 10^{-19}$ |
| TRANS | PHOS | 266 | 0.241723 |
| PROT | ACET | 133 | $8.53 \times 10^{-05}$ |
| PROT | PHOS | 117 | 0.925481 |
| ACET | PHOS | 89 | 0.420549 |

**B**

| Regulatory mechanisms | | Gene Overlap | p-value |
|---|---|---|---|
| TRANS | PROT | 153 | 0.010941 |
| TRANS | ACET | 157 | 0.001552 |
| TRANS | PHOS | 61 | 0.931005 |
| PROT | ACET | 69 | 0.925509 |
| PROT | PHOS | 42 | 0.291789 |
| ACET | PHOS | 34 | 0.860463 |

**C**

| Total regulators | Percentage among those regulated |
|---|---|
| 2 or more | 47.8% |
| 3 or more | 8.7% |
| All 4 | 0.08% |

**S. Table 1.** Overlap between targets of various mechanisms - transcription (TRANS), post-transcription (PROT), acetylation (ACET), phosphorylation (PHOS) in yeast. This reveals low overlap between targets of regulation by phosphorylation and other mechanisms. **A.** Overlap between target reactions **B.** Overlap between target genes. **C.** Percentage of reactions regulated by multiple mechanisms. Overall, 69% of the gene-associated reactions in the model were regulated; among those regulated, 47.8% were regulated by more than one mechanism.

**S. Table 2.** Essential reactions regulated by acetylation in yeast (Spreadsheet file)

**S. Table 3.** Top 50 reactions sorted based on topological connectivity in yeast (Spreadsheet file)

**S. Table 4.** Top 50 reactions with maximum reaction flux regulated by phosphorylation in yeast (Spreadsheet file)

| Model | Yeast 7 (default) | | Yeast 7.6 | |
|---|---|---|---|---|
| p-value | ANOVA | Kruskal-Wallis | ANOVA | Kruskal-Wallis |
| Growth rate | $2.07 \times 10^{-41}$ | $1.24 \times 10^{-29}$ | $7.82 \times 10^{-43}$ | $1.37 \times 10^{-47}$ |
| Closeness | $3.33 \times 10^{-48}$ | $1.74 \times 10^{-55}$ | $1.66 \times 10^{-39}$ | $3.61 \times 10^{-51}$ |
| Vmax (without max. biomass) | $5.53 \times 10^{-26}$ | $9.25 \times 10^{-13}$ | $6.97 \times 10^{-8}$ | $1.30 \times 10^{-11}$ |

**S. Table 5.** Robustness of the results comparing the difference in distribution of properties between targets of various regulatory mechanisms using the Yeast 7.6 model. Significance of results using the non-parametric Kruskal-Wallis test is also shown. The p-values for the key reaction features shown in Figure 1 using the Yeast 7 model is provided as comparison. All p-values are significant at FDR < 0.01 using both Bonferroni adjustment and Benjamin-Hochberg multiple hypothesis correction.

| Model | All genes (default) | | All expressed genes | |
|---|---|---|---|---|
| p-value | ANOVA | Kruskal-Wallis | ANOVA | Kruskal-Wallis |
| Growth rate | $2.07 \times 10^{-41}$ | $1.24 \times 10^{-29}$ | $3.3 \times 10^{-41}$ | $2.1 \times 10^{-29}$ |
| Closeness | $3.33 \times 10^{-48}$ | $1.74 \times 10^{-55}$ | $2.2 \times 10^{-49}$ | $3.5 \times 10^{-56}$ |
| Vmax | $1.59 \times 10^{-26}$ | $2.51 \times 10^{-21}$ | $8.1 \times 10^{-27}$ | $1.3 \times 10^{-21}$ |

**S. Table 6.** Robustness of the results after removing genes that are not-expressed (i.e. not detected in RNA-seq data) in both exponential and stationary phase cultures. The p-values reported in Figure 1 using all the metabolic genes in the Yeast 7 model is provided as comparison.

| Model | Murphy *et al* (default) | | Weinert *et al* | |
|---|---|---|---|---|
| p-value | ANOVA | Kruskal-Wallis | ANOVA | Kruskal-Wallis |
| Growth rate | $2.07 \times 10^{-41}$ | $1.24 \times 10^{-29}$ | $1.59 \times 10^{-37}$ | $2.63 \times 10^{-33}$ |
| Closeness | $3.33 \times 10^{-48}$ | $1.74 \times 10^{-55}$ | $3.10 \times 10^{-44}$ | $1.62 \times 10^{-48}$ |
| Vmax | $1.59 \times 10^{-26}$ | $2.51 \times 10^{-21}$ | $1.71 \times 10^{-22}$ | $1.94 \times 10^{-19}$ |

**S. Table 7.** Comparison of results using proteomics data from Weinert *et al* instead of Murphy *et al*. The ANOVA p-value comparing the means are provided. The p-values reported in Figure 1 using Murphy *et al* data is provided as comparison.

| Fold change | 2 (default) | 1.5 | 3 | 4 |
|---|---|---|---|---|
| Growth rate | $2.07 X 10^{-41}$ | $3.72 X 10^{-35}$ | $1.12 X 10^{-34}$ | $5.82 X 10^{-26}$ |
| Closeness | $3.33 X 10^{-48}$ | $2.09 X 10^{-47}$ | $1.13 X 10^{-32}$ | $6.95 X 10^{-20}$ |
| Vmax (with max. biomass) | $1.59 X 10^{-26}$ | $6.97 X 10^{-24}$ | $1.26 X 10^{-23}$ | $1.01 X 10^{-19}$ |

| Top Percentile | 25 (default) | 50 | 15 | 5 |
|---|---|---|---|---|
| Growth rate | $2.07 X 10^{-41}$ | $1.42 X 10^{-42}$ | $2.49 X 10^{-39}$ | $2.71 X 10^{-40}$ |
| Closeness | $3.33 X 10^{-48}$ | $1.86 X 10^{-41}$ | $7.15 X 10^{-55}$ | $9.66 X 10^{-48}$ |
| Vmax (with max. biomass) | $1.59 X 10^{-26}$ | $6.07 X 10^{-18}$ | $5.66 X 10^{-31}$ | $8.98 X 10^{-36}$ |

**S. Table 8.** Comparison of thresholds used for identifying differentially expressed genes and proteins. These show that our results are robust to the thresholds for identifying the targets of various regulatory mechanisms. The ANOVA p-value comparing the means are provided. Note that the first table uses fold change thresholds for transcriptomics, acetylation and phospho-proteomics data alone. Since the proteomics data uses a percentile cut off, the robustness analysis for this data was performed separately.

| Threshold for unconstrained reactions | ANOVA p-value for Vmax |
|---|---|
| 100 | $5.07 x 10^{-31}$ |
| 200 | $1.08 x 10^{-57}$ |
| 300 | $5.27 x 10^{-50}$ |
| 400 | $3.53 x 10^{-27}$ |
| 500 | $2.24 x 10^{-25}$ |
| 600 | $2.24 x 10^{-25}$ |
| 700 | $2.24 x 10^{-25}$ |
| 800 | $1.59 x 10^{-26}$ |
| 900 | $1.59 x 10^{-26}$ |
| 1000 | $4.68 x 10^{-150}$ |

**S. Table 9.** Comparison of thresholds used for identifying unconstrained reactions from FVA. Reactions with maximal flux above the threshold listed in the table were assumed to be unconstrained and were excluded from the analysis, as they are likely due to thermodynamically infeasible internal cycles. The ANOVA p-value comparing the means of the maximum flux through the target reactions of different regulatory mechanisms is provided. The default value (900 mmol/gDW/hr) for eliminating unconstrained reactions is highlighted and was used for all analyses. These show that our results are robust to the thresholds for identifying unconstrained reactions.

| Regulation | *E. coli* | S. cerevisiae |
|---|---|---|
| Transcription | 469 | 468 |
| Post-transcription/Proteomic | 372 | 266 |
| Acetylation | 460 | 265 |
| Phosphorylation | 17 | 133 |

**S. Table 10.** Comparison of total genes regulated by each process in *E. coli* with *S. cerevisiae* shows that phosphorylation plays a relatively minor role in *E. coli* metabolic regulation during stationary phase.

| Regulatory mechanisms | | p-value | Reaction Overlap |
|---|---|---|---|
| TRANS | PROT | $3.77 \times 10^{-23}$ | 590 |
| ACET | TRANS | 0.042022 | 442 |
| ACET | PROT | 0.192068 | 379 |
| ACET | PHOS | $5.60 \times 10^{-11}$ | 28 |
| PHOS | TRANS | 0.004853 | 22 |
| PHOS | PROT | 0.95148 | 9 |

**S. Table 11.** Overlap between targets of various mechanisms in *E. coli* - transcription (TRANS), post-transcription (PROT), acetylation (ACET), phosphorylation (PHOS).

**S. Table 12.** Dataset containing phosphorylation predictions in the cell cycle data (Spreadsheet file).

**S. Table 13.** Dataset containing acetylation predictions in the cell cycle data (Spreadsheet file).

**S Table 14.** Hypergeometric test results for additional lysine deacetylase inhibitors from the Scholz *et al.* study. The number of unique acetylated genes for each group are displayed in parentheses. Within the table, the number of overlapping genes between each phase and drug is shown, along with the upper tail p-value of the hypergeometric test. (Spreadsheet file).

**S. Table 15.** Raw dataset containing all yeast genes and associated reactions, the corresponding regulators, and the reaction properties (Spreadsheet file).

**S. Table 16.** Raw dataset containing all E. coli genes and associated reactions, the corresponding regulators, and the reaction properties (Spreadsheet file).

**S. Table 17.** Raw dataset containing all human genes and associated reactions, the corresponding regulators, and the reaction properties (Spreadsheet file).

**S. Table 18.** Raw dataset containing the phosphoproteomics data from the cell cycle G0 phase for training the model (Spreadsheet file).

**S. Table 19.** Final normalized dataset used for training the CAROM model (Spreadsheet file).

**S. Table 20**. Overlap between genes that were predicted by CAROM and those found to be differentially acetylated for the four pan-lysine deacetylase inhibitors that we analyzed from the Scholz *et al.* study. (Spreadsheet file).
